# Herding cats: predicting immunogenicity from heterogeneous clinical trials data

**DOI:** 10.1101/2025.10.14.682375

**Authors:** Pawel Dudzic, Konrad Krawczyk

**Affiliations:** NaturalAntibody & Friends

## Abstract

Antibodies represent the largest and fastest growing class of biologic therapeutics, yet forecasting their clinical performance, particularly immunogenicity, remains a major hurdle in drug development. Despite hundreds of antibody-based drugs progressing through clinical pipelines, systematic integration of their clinical outcomes has been limited by fragmented and heterogeneous data. Here, we present the Therapeutic Antibody Database, a comprehensive and curated resource that links therapeutic antibodies to clinical trial outcomes, with a dedicated focus on immunogenicity. Our dataset is sourced from approximately 11,500 anti-drug antibody (ADA) measurements across diverse molecules and indications, offering an unprecedented view into the clinical manifestation of immune responses to biologics. In order to evaluate the main drivers of ADA, we evaluate gathered immunogenicity incidence and prevalence data against various therapeutic descriptors which includes sequence, structure and contextual features related to therapeutics. We find that most tools have very poor performance, and we pinpoint the causes of it, demonstrating the need for systems immunology approaches incorporating clinical metadata beyond biochemical properties of the molecules alone.

## Introduction

Therapeutic antibodies have revolutionized modern medicine, offering targeted treatments for a wide range of diseases (Walsh and Walsh 2022). With antibodies being the majority of blockbuster drugs on the market, immunogenicity remains a major problem in this class of therapeutics (Hansel et al. 2010; Carter and Quarmby 2024). This unwanted patient reaction, in which anti-drug antibodies (ADA) are being created, can render the drug ineffective, alter its pharmacokinetics, or lead to severe adverse effects. In some cases it can even lead to whole therapeutic programs being shut down (Carter and Quarmby 2024). Immunogenicity is a multifaceted problem with factors broadly categorized into product and patient-related characteristics, where in all cases the presence of ADA is associated with B-cell or T-cell pathways (Sun, Qian, and Zhang 2024).

The first therapeutic monoclonal antibodies were obtained by immunizing mice, yielding antibodies fully “murine” in structure but highly specific for a given antigen (e.g. muromonab, approved in 1986). It quickly became clear that administering a protein of a xeno origin to patients elicits an unwanted immune response, significantly limiting treatment efficacy. Consequently, sequence-modification strategies were introduced - first chimerization, in which murine constant regions were replaced with their human counterparts, and later humanization, in which the antigen-recognition regions (CDRs - complementarity-determining regions, or in some protocols SDRs - specificity-determining residues) were grafted onto a “human” framework (Safdari et al. 2013; Jones et al. 1986). The underlying assumption was the intuition that the more “human” the protein translates the lower the risk of immunogenicity (Baker et al. 2020).

Today, owing to display technologies (phage display, mammalian display) and transgenic mice carrying human immune system genes, fully human sequences can be generated (Lonberg 2008). Nevertheless, anti-drug antibodies are still observed. The benefit of such ‘humanization’ methods has been previously challenged (Clark 2000); for example Abhinandan and Martin (Abhinandan and Martin 2007), compared human antibody sequence content, or “humanness”, with measured immunogenicity and found no correlation. Moreover, in practice many companies still apply various humanization protocols not always grounded in a deep mechanistic understanding of immunogenicity, rather focusing on human sequence content with some resort to critical residues that might affect binding (e.g. Vernier zone residues). Paradoxically, as we show later, there are murine antibodies that are well tolerated clinically and fully human antibodies with high immunogenicity, indicating that this is a much more complex problem that cannot be easily reduced to simple ‘human vs murine content’ alone.

When a protein therapeutic is administered to the patient, it gets internalized by antigen-presenting cells (APC) such as macrophages, dendritic cells, or B-cells. The protein is further split into peptides with proteases in lysosomes, so shorter fragments of the amino acid sequence can bind to the MHC II complex and get exposed on the surface of APC for the T-cell for recognition. The activation of a T-cell requires not only epitope recognition on MHC but also costimulatory signals. For example, in the presence of PAMP and DAMP molecules, dendritic cells get activated and present molecules such as CD80 or CD86 on their surface, which interact with CD28 on the T-cell. Costimulatory signals induce the presence of CTLA4 molecules on the surface of the T-cell. It has an inhibitory effect, but it also has a higher affinity to CD80/CD86 therefore it “frees” CD28. Further activation and the release of the cytokines are therefore a result of a balance between inhibitory and costimulatory signals, which in turn activate affinity maturation of B-cells which start to produce ADA with high affinity to the protein. To understand the importance of this process, it is necessary to explore the immunogenic T-cell epitopes within the sequence, in the therapeutic context, such as the mechanism of action or patient disease, as it is done in this paper.

In the B-cell dependent pathway, the primary reason for immunogenicity is when the therapeutic protein encounters pre-existing B-cell epitopes (Carter and Quarmby 2024). As each patient possesses a unique set of B-cell receptors, this inherently difficult problem is further complicated by the presence of opportunistic epitopes, emergent after the therapeutic protein aggregation. Apart from aggregation propensity, previous studies associated the protein charge or charged patches with immunogenicity (Carter and Quarmby 2024), because of increased therapeutic internalization, which increases the exposition for the T-cell receptors.

As biologics become increasingly central to patient care, comprehensive and easily accessible databases are essential for guiding research and clinical decision-making. While several existing antibody-focused repositories provide valuable information, they often concentrate solely on variable regions, exclude essential clinical data on immunogenicity, lack details on post-translational modifications (PTMs) or are very small (e.g. 217 unannotated points (Marks et al. 2021)).

To address these critical gaps, we present the Therapeutic Antibody Database (naturalantibody.com/therapeutic-antibody-database/), a fully integrated database encompassing not only complete antibody sequences, with both constant and variable regions, but also extensive PTM profiles, clinical trial linkage, and a robust collection of manually curated immunogenicity scores (ca. 11,500). With its breadth and depth, this comprehensive dataset stands as the largest current resource of its kind, enabling unprecedented stratification of risk factors related to product characteristics and patient attributes such as the disease, and ultimately driving improved safety and efficacy in therapeutic antibody development. Using this curated resource, we explore various sequence, structure and context features that might influence the emergence of ADA. Notably: humanness scores of variable regions, the presence of immunogenic T-cell epitopes, protein electrostatics charged surface patches descriptors, and therapeutic context, such as the mechanism of action or patient disease.

## Results

### Database statistics

The therapeutic database contains full sequence information, post translational modifications and target annotations for 1,109 therapeutics, gathered chiefly from the INN lists published by World Health Organization and manual searches on clinical trials data. The therapeutics were assigned to 117 distinct formats, categorized into 5 groups (Figure 1A). We have identified 438 unique targets which were augmented with UniProt data.

**Figure 1.**
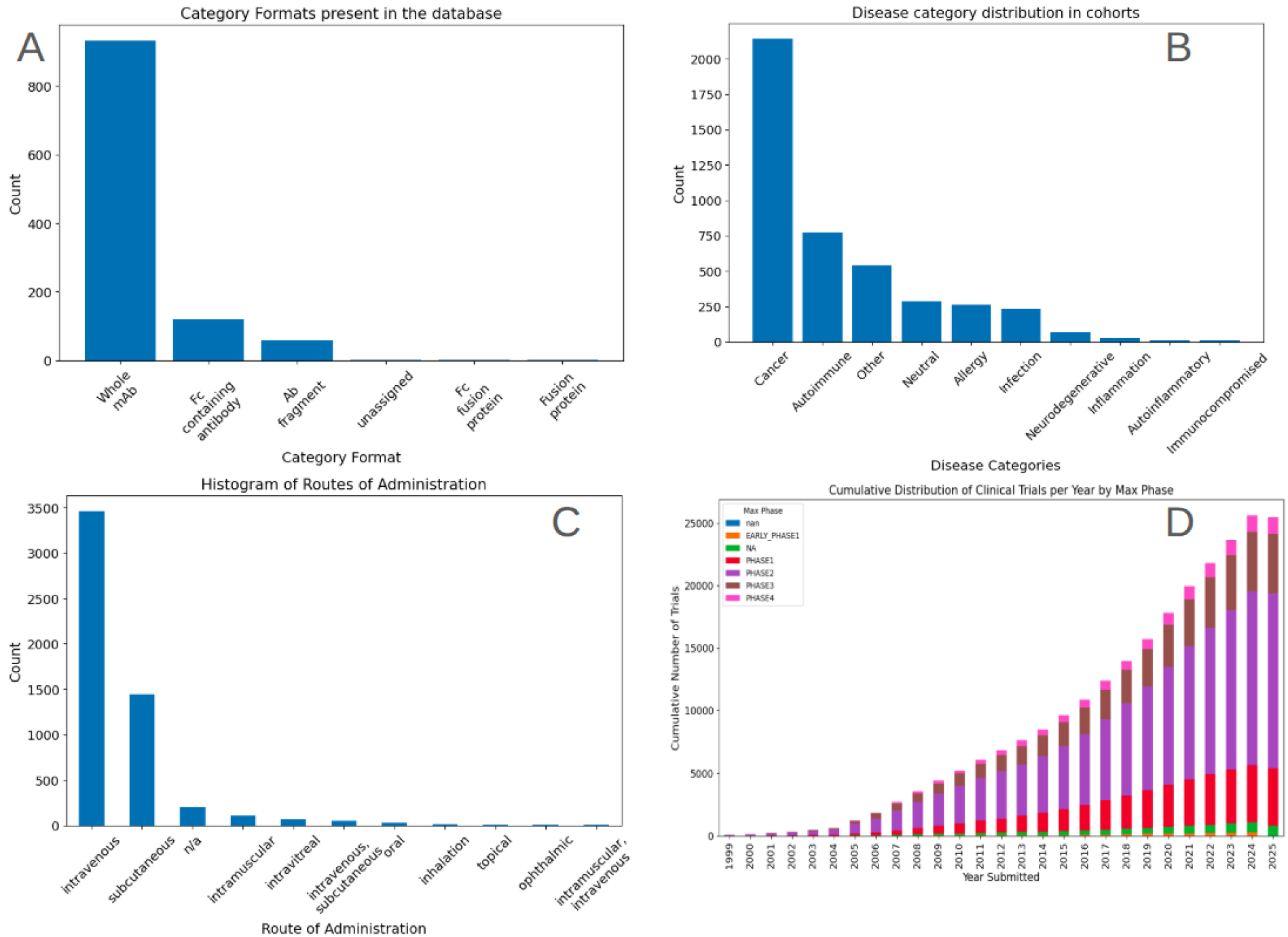
Database overview. Therapeutic format categories present in the database. (A). Distribution of disease categories assigned to cohorts (B). Distribution of routes of administration for all interventions (C). Cumulative number of antibody therapeutic related clinical trials (note that we do not have full data for 2025 at the time of writing) (D).

For all of the therapeutics from the database, we have searched clinicaltrials.gov using the names and synonyms. This yielded therapeutics-related clinical trials, measuring distinct outcomes. We have identified clinical trials in which immunogenicity was measured using our in-house semantic search engine. This yielded 1,120 clinical trials measuring immunogenicity data in 4,306 cohorts, with the overall number of identified measurements equal to 11,575, which are all present in the database. The measurements cover 300 distinct therapeutics. There are a total of 277 unique diseases assigned to all cohorts, which we grouped into 10 categories (Figure 1B). We have identified a total of 6,961 cohort interventions and assigned the dosages and routes of administration to them (Figure 1C). The number of therapeutic antibody related clinical trials rises steadily (Figure 1D), indicating that the value of resources such as this one will only gather in volume and thus value with time.

Since the popularity of the targeted molecules directly corresponds to clinical relevance, providing the link between the target and the disease, we have counted the occurrences of popular targets of therapeutics. Targets with 10 or more therapeutics aiming at them are presented in Table

We have identified several obstacles during data curation, which rendered comparisons of the measurements between the clinical trials challenging. For instance, sometimes there was ambiguity in reporting the number of ADA patients being positive or negative, or uncertainty when the measurements were performed (e.g. before or after the drug administration). Critically, we identified a need to categorize the measurements according to their relationship to pre-drug administration status (e.g. whether measurement was performed on baseline positive or negative only). To address this, we have harmonized the measurements, as described in the annotation pipeline, to produce the subset datasets, representing incidence, prevalence, and pre-existing immunogenicity (Figure 2), consisting of 1651, 3859, and 1678 records, respectively.

**Figure 2.**
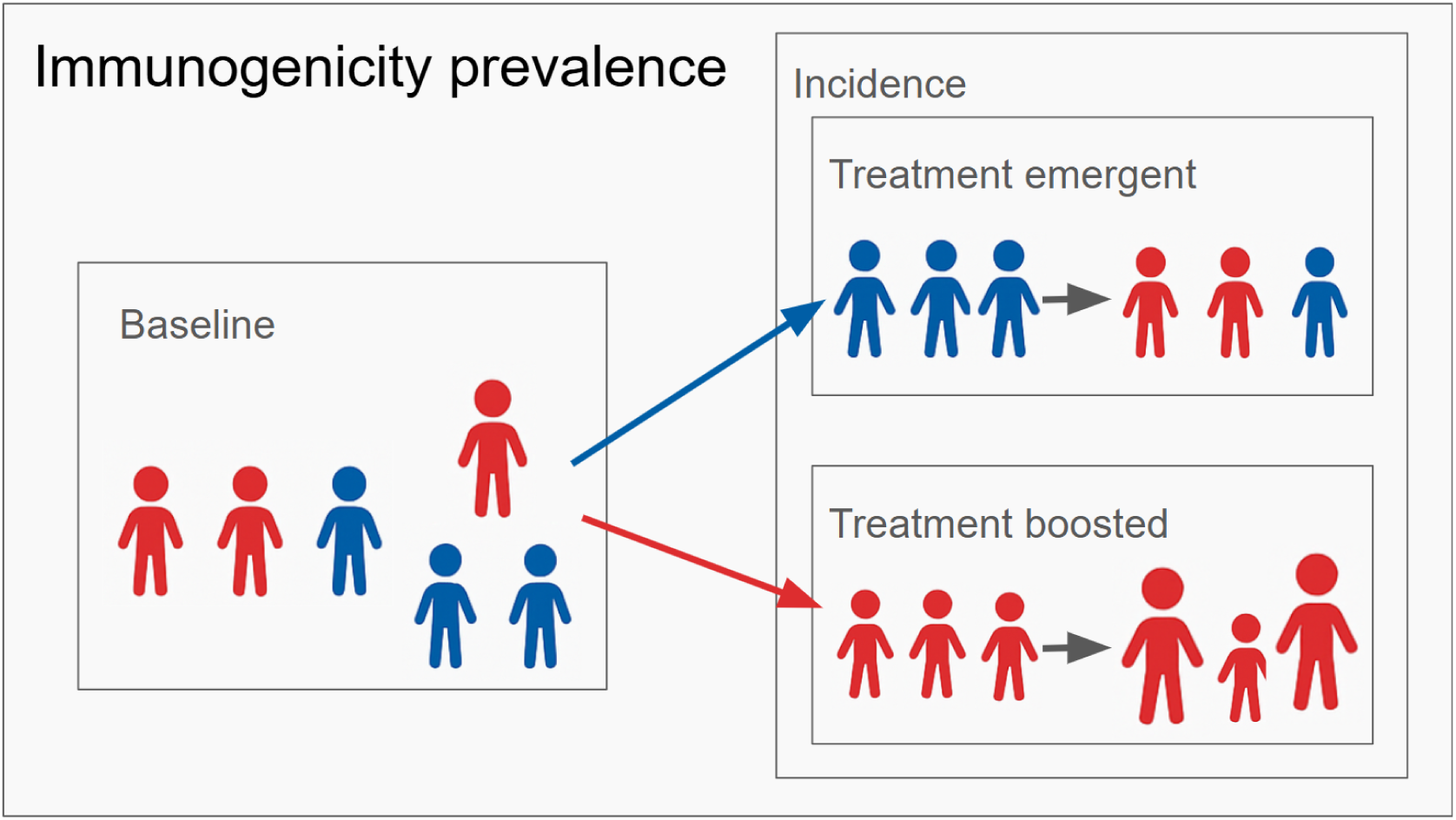
Different types of immunogenicity measurements. Blue color represents ADA negative patients and red color - ADA positive. Increase in ‘person’ icon size indicates increase in ADA titers in treatment boosted immunogenicity. When no stratification is made in the measurement with respect to baseline status, we treat the values as ADA prevalence. Baseline measurements are performed before the drug administration. Treatment emergent ADA is the number of positive patients that were negative at baseline. Treatment boosted ADA is the number of baseline positive patients where the ADA titers increased (at least 4-fold increase in titers). Incidence consists of both: boosted and emergent. Prevalence is incidence with baseline.

We define immunogenicity incidence as the presence of treatment-emergent or boosted ADA so that only relative measurements are considered, performed after the drug administration. Treatment emergent measurement is defined as the number of ADA-positive participants (or percentage of participants) who were ADA-negative at baseline (before the drug administration). Similarly, treatment boosted measurements are defined as positive post-baseline participants, who were also positive before the drug administration, and there was a significant change in ADA titer levels (usually 4-fold). Immunogenicity prevalence is defined as the measurements of all patients without the stratification on the baseline results - the incidence, but together with baseline positive measurements. Finally the baseline dataset was constructed to investigate the pre-existing immunogenicity of the therapeutics. To build this subset, we gathered all of the measurements from placebo groups (675 measurements) and all groups in which the ADA was measured before the drug administration (1097 measurements).

When considering immunogenicity data, one should consider the significant evolution in assay technology and sensitivity. Early clinical trials data were often inconsistent making them incomparable to results generated from modern standardized trials (Bawa et al. 2019). To minimize the problem of varying titer sensitivity, for the following exploratory data analysis, only the measurements from studies completed in 2016 or after were considered.

### Therapeutic molecule features are insufficient for immunogenicity prediction

#### Mouse sequence contents is not the chief determinant of immunogenicity

To determine sequence characteristics responsible for ADA formation we contrasted ADA to various humanness scores, estimated T-cell immunogenicity and biophysical sequence descriptors. To explore the relationship between the mouse sequence contents and the immunogenicity in our dataset, we first visualized the prevalence and incidence for each therapeutic type (Figure 3 A and B), then we visualized the humanness of therapeutics variable regions to the immunogenicity prevalence and incidence (Figure 3 C and D). We define humanness score as a sum of alignment scores of the therapeutic variable regions to the closest V and J germline sequences, as assigned by RIOT annotations (Paweł Dudzic et al. 2024).

**Figure 3.**
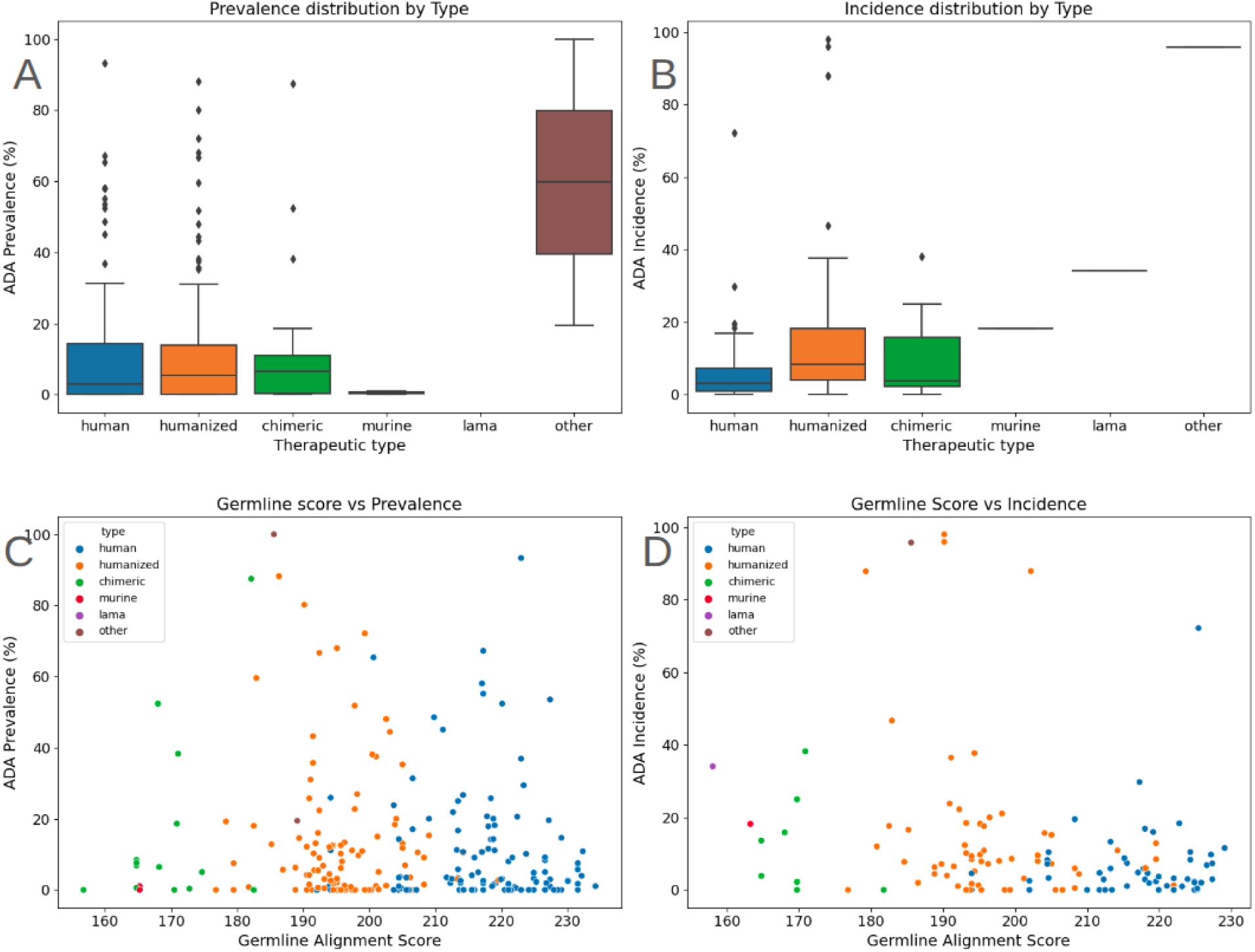
Distribution of immunogenicity in different groups of therapeutics. (A) Prevalence. (B) Incidence. Human germline similarity compared to immunogenicity measurements for different types of therapeutics. (C) Prevalence. (D) Incidence.

While the average immunogenicity incidence (3D) is the lowest for human therapeutics, we still observe the variability: there are non-human like therapeutics with low immunogenicity and therapeutics close to human germlines for which there exists a number of clinical outcomes where immunogenicity is noted.

Immunogenicity prevalence (3C) is more difficult to interpret - on one hand we have much more data, on the other, the prevalence captures both: pre-existing immunogenicity and treatment induced one. Here the difference between the human and humanized therapeutics is blurred. While the measurements could be easily separated by the distance to the closest human germlines, there is no clear gain in terms of observed ADA scores. The low prevalence of anti-drug antibodies (ADA) in murine therapeutics is attributed to the limited available data.

We further explored the existing tools used to assess the humanness scores. We scored variable regions of all therapeutics in the database using BioPhi OASis (Prihoda et al. 2022), Hu-mAb (Marks et al. 2021), Shab (Abhinandan and Martin 2007) and T20 (Gao et al. 2013) scores. In most cases, each of the chains was scored separately and the average score from both chains was used for comparison to ADA values. As Hu-mAb produces several scores for each chain (one similarity score to each V gene subgroup), we selected the highest score for each chain, and from those best scores the average was calculated. None of the metrics showed statistically significant correlations with either ADA prevalence (Figure 4) nor incidence (Figure 5) (p>0.05 in all cases with the absolute value of Pearson correlation coefficient less than 0.19).

**Figure 4.**
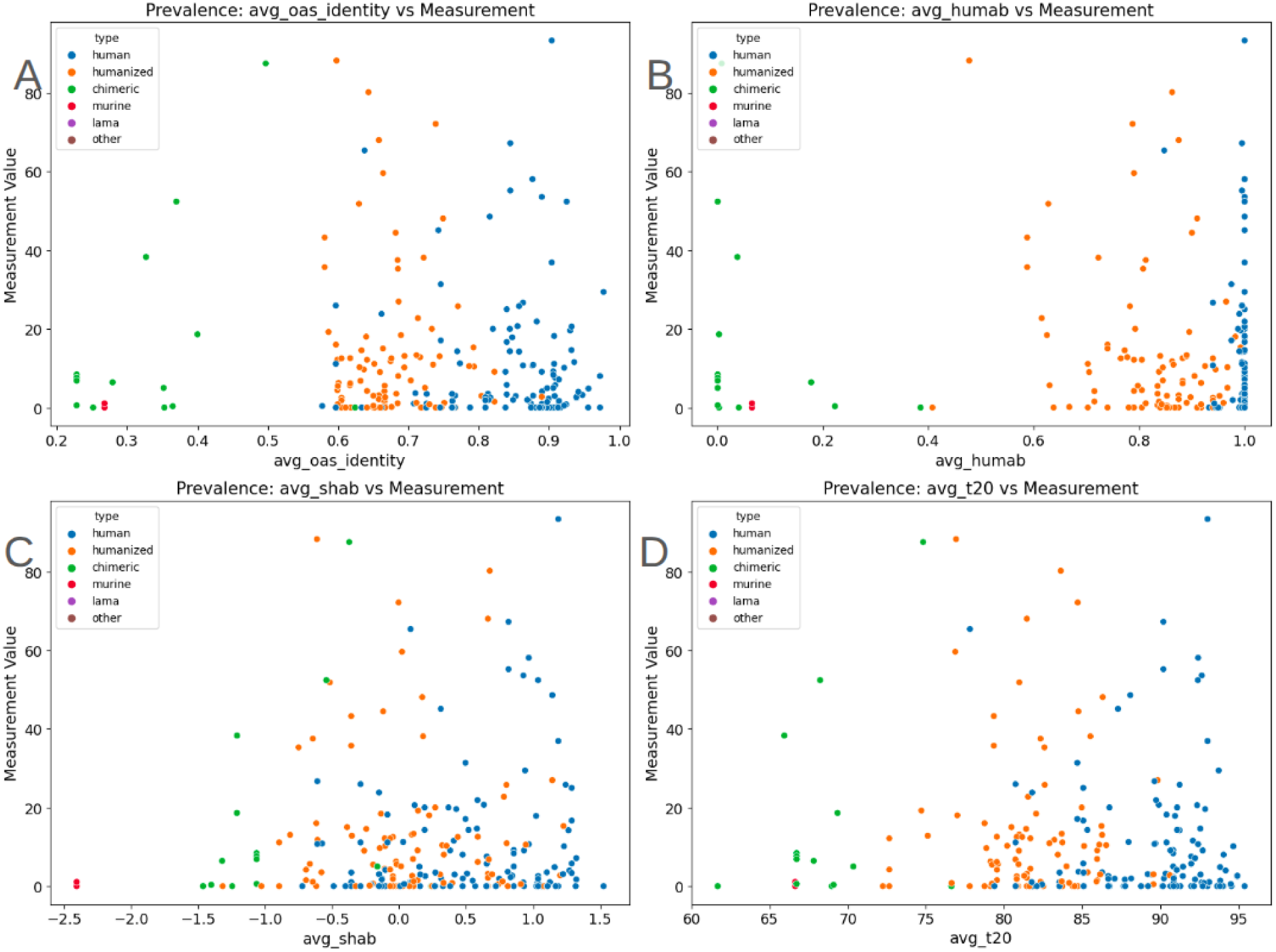
Humanness scores against immunogenicity prevalence for OAS identity (A), Hu-mAb (B), shab (C) and t20 (D). While scores separate distinct antibody types based on human sequence content, none of the humanness scores shows statistically significant correlation to immunogenicity prevalence.

**Figure 5:**
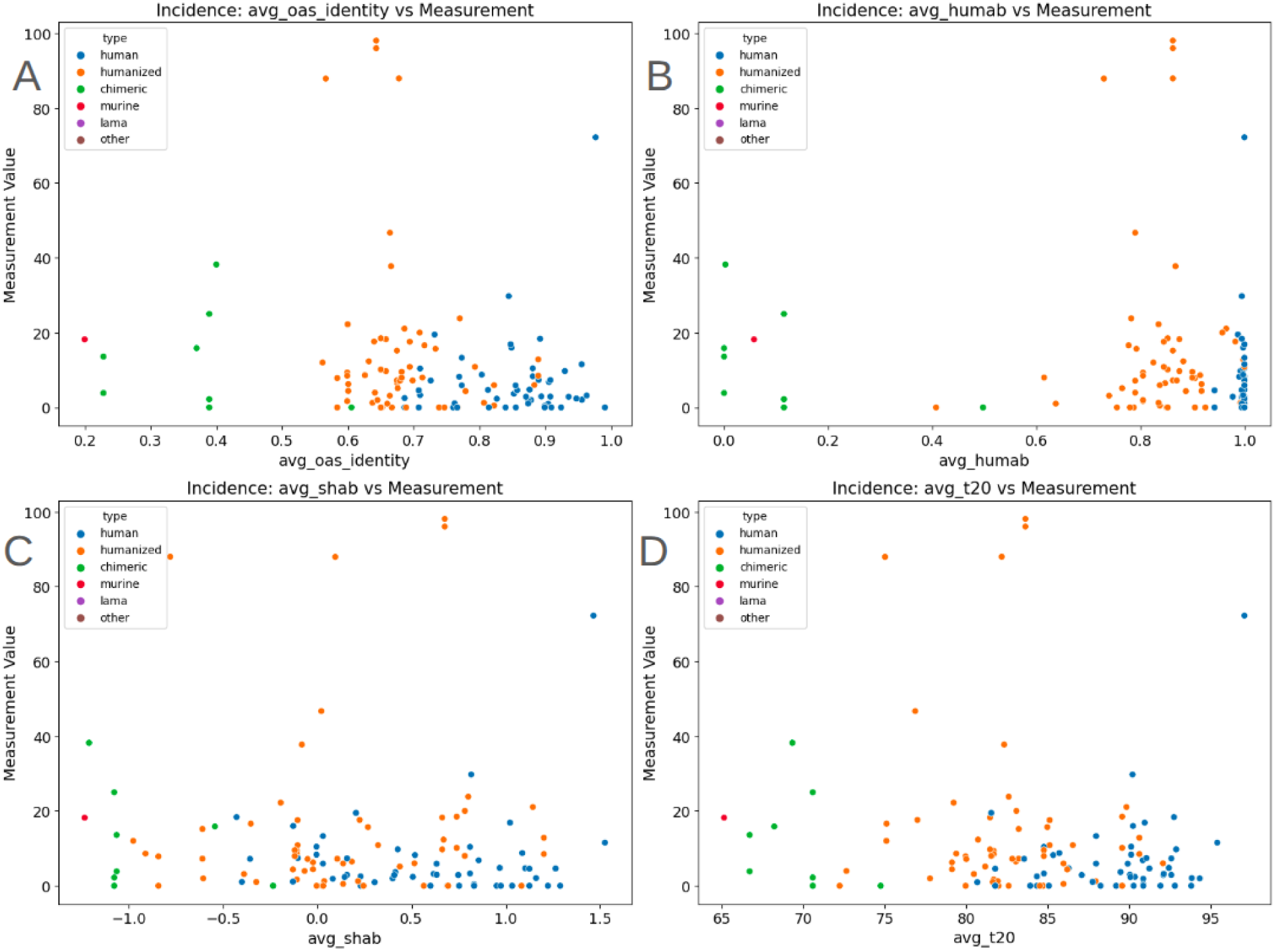
Humanness scores compared to immunogenicity incidence for OAS identity (A), Hu-mAb (B), shab (C) and t20 (D). While there is a visible separation of distinct antibody types, none of the humanness scores shows statistically significant correlation to immunogenicity incidence.

Given the above results, it appears that humanness does not have strong predictive power of ADAs, as the simple germline alignment score segments different therapeutic types. The results are consistent with other work (Hu, Cohen, and Swanson 2023). However, while we did not find evidence supporting the use of humanness as a direct reflection of immunogenicity, there are works which link naturalness of the proteins to better developability scores (Bachas et al. 2022). Therefore naturalness/humanness of sequences should not be omitted but also should not be treated as a single factor that will predict/reduce immunogenicity.

#### In-silico predicted immunogenic T-cell epitopes do not determine the immunogenicity

Identifying putative immunogenic T-cell epitopes is a crucial starting point for assessing a biologic’s immunogenicity risk, yet their mere presence does not consistently translate into clinical antidrug antibody formation (A. S. De Groot, Knopp, and Martin 2005). This discrepancy underscores the multifaceted nature of immune recognition, including how antigen processing and presentation can mask or limit epitope availability. As a result, future work should focus on examining these intracellular and extracellular pathways to gain clearer insight into why certain predicted epitopes remain clinically silent.

In order to compare the presence of immunogenic T-cell epitopes to clinical immunogenicity, we predicted T-cell epitopes in variable regions. All overlapping peptides from variable regions were scored with CD4Episcore (Dhanda et al. 2018). From the pool of peptides, we removed ones abundant in our PairedNGS dataset (Pawel Dudzic et al. 2025) of natural paired human heavy and light chains. The final score of the therapeutic was calculated by summing the scores from all of the potential epitopes. The results for prevalence and incidence are presented on Figure 6.

**Figure 6.**
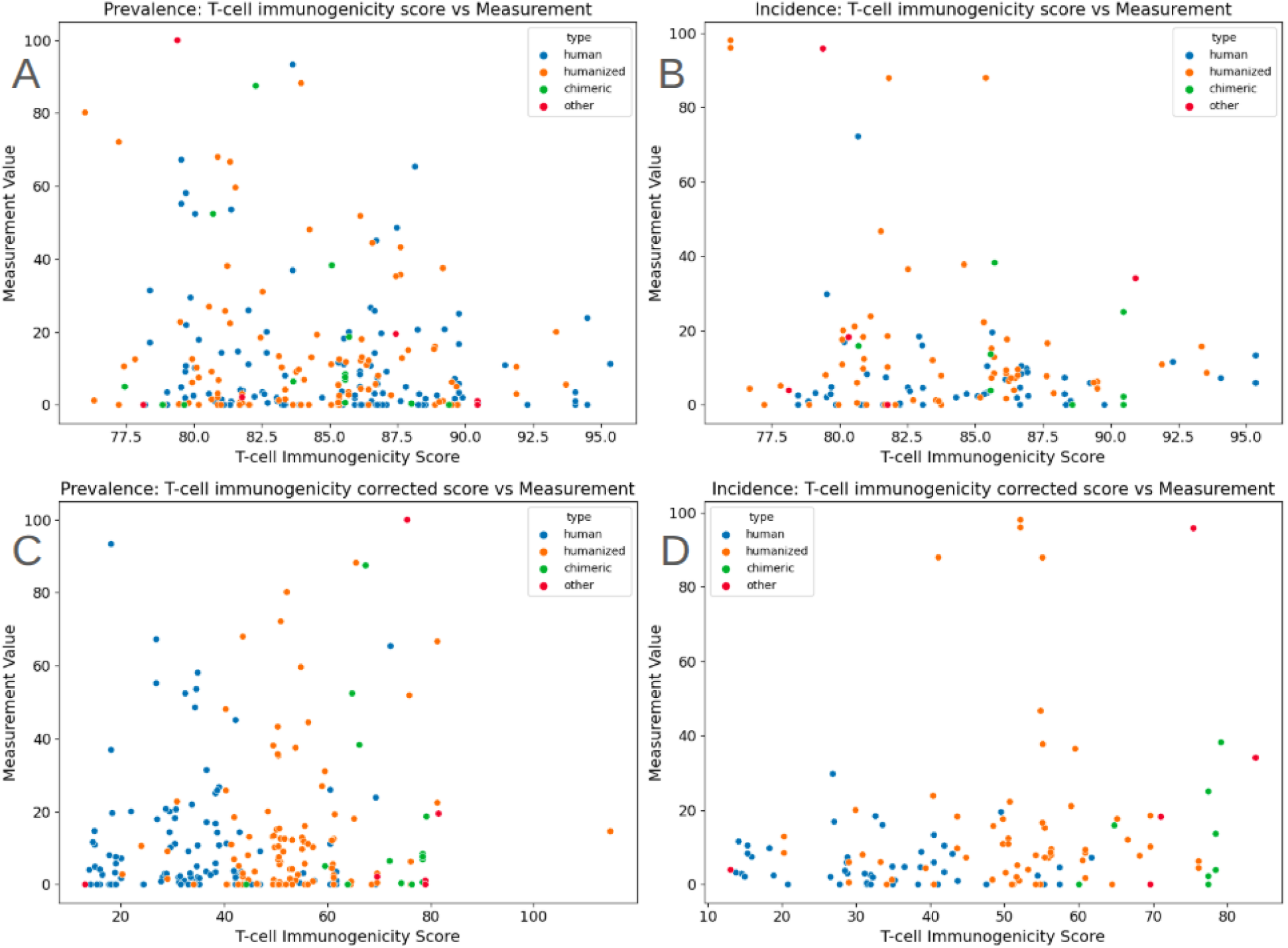
The CD4 activation score of therapeutics against clinical measurements of immunogenicity. Considering all predicted immunogenic epitopes, there is no clear separation between therapeutic types (A, B). Ignoring well tolerated peptides renders better therapeutic type separation by the score, yet does not improve the correlations (C, D).

The results were stratified by the therapeutic type. We noted that from the sequence point of view, more human-like therapeutics are not associated with reduced T-cell immunogenic score of a therapeutic (there is no clear distinction between different types of therapeutics, Figure 6 A, B) and does not correlate statistically significant with ADA prevalence nor incidence. After removing peptides abundant in human sequencing studies, despite expected better separation of therapeutic types, we have not observed any improvement in the correlations (Figure 6 C, D). This discrepancy suggests that factors beyond unpopular epitope presence - most notably antigen processing - may obscure or diminish the immunogenic potential of therapeutic proteins. The results are consistent with a previous study (Tsai et al. 2024).

While we haven’t observed any correlations, the results should be interpreted with caution - most of the therapeutics had probably undergone some kind of deimmunization or T-cell epitope removal. Therefore proposed in-silico protocol should be validated experimentally using MAPP assays to determine true epitopes capable of MHC II binding. Furthermore, investigations targeting the mechanisms of epitope display and proteolytic processing could provide crucial insights into why certain predicted epitopes fail to elicit antidrug antibodies and ultimately help refine risk assessments for novel biologics.

#### No free lunch: sequence and biophysical features are incapable of immunogenicity prediction using simple machine learning models

We have explored various basic biophysical variable region features derived either from the whole variable fragment (including heavy and light chains), as well as features derived from surface exposed residues in the variable region, such as surface aggregation propensity (SAP) (Black and Mould 1991). As the T-cell dependent pathway is primarily driven by the sequence features, the B-cell pathway, caused by the presence of pre-existing and the opportunistic B-cell epitopes, emergent after therapeutic aggregation, is associated with structural features of proteins (Moussa et al. 2016; Ratanji et al. 2014). Furthermore, aggregation is linked to Toll-like receptors activation and increased uptake of therapeutic aggregates by APC. To explore this pathway of immunogenicity, we calculated structural descriptors for electrostatic surface potentials integrals and aggregation propensities for surface exposed residues, for the ABodyBuilder3 (Kenlay et al. 2024) generated variable fragments structures (see methods).

We used machine learning to determine features related to ADA presence. The dataset was enriched with a multitude of therapeutic features and context variables. Therapeutic features included humanness scores, T-cell immunogenicity scores, therapeutic mode of action determined after its target interaction type, electrostatics surface potential integrals for frameworks and CDRs, and surface aggregation propensities scores calculated for each variable fragment region, basing on surface exposed residues. Context dependent features included cohort disease and the route of administration. Each data point in prevalence dataset was flagged as immunogenic when the immunogenicity measurement was above 10%.

Using this dataset, we have trained a random forest classifier in a two step process: at first the model was trained on all of the available features. Then scikit-learn random forest built in feature importance was used to select only the most relevant features for the second iteration of model training. The threshold of 0.02 mean accuracy decrease was used to filter the features. With the features selected, the model was retrained and evaluated on the test set of therapeutics not present in training. While achieving moderate predictive capabilities (AUC = 0.72), we compared a series of metrics of the model to the predictions of the model trained on the same dataset, where the labels were randomly permuted (Figure 7), revealing very poor performance of the model. Furthermore, the permutation based feature importance showed only a handful of relevant features sensitive to the data split. We concluded that the model was unable to generate reliable predictions.

**Figure 7.**
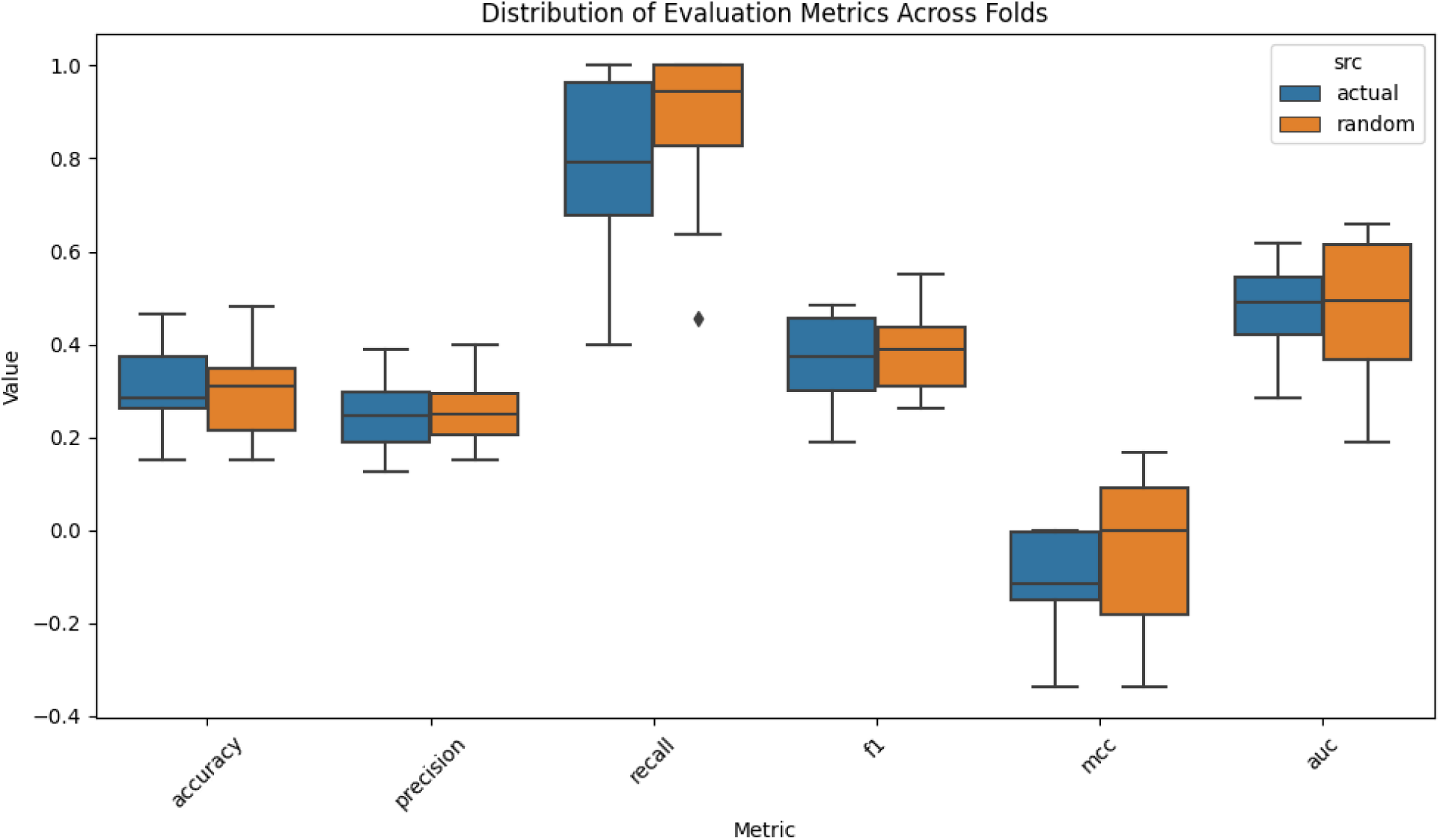
The random forest performance metrics. The model was evaluated on the therapeutics not present in the training data. The metrics were compared to the same test data where the labels were randomly permuted which maintained the data distribution.

### Drowning in the noise: immunogenicity measurements are highly variable

To elucidate the poor performance of the predictive model we further explored the variability of the immunogenicity measurements for individual therapeutics as exemplified by Adalimumab and Atezolizumab (Figure 8), which is present in both prevalence and incidence measurements (Figure 9 A and B respectively). Altogether the issue lies in large clinical context dependence of the therapeutic. In some extreme cases, a single molecule can have immunogenicity measurements covering the entire 0%-100% scale, if the clinical context is changed. As an example we plot the measurements of immunogenicity for the two of therapeutics to depict the issue in Figure 8. Note that both Adalimumab and Atezolizumab have a very wide spread of immunogenicity - despite the ‘molecular features’ arguably being the same for each measurement. Such wide-ranging variability is reflected when we look at the totality of therapeutics as well (Figure 9).

**Figure 8.**
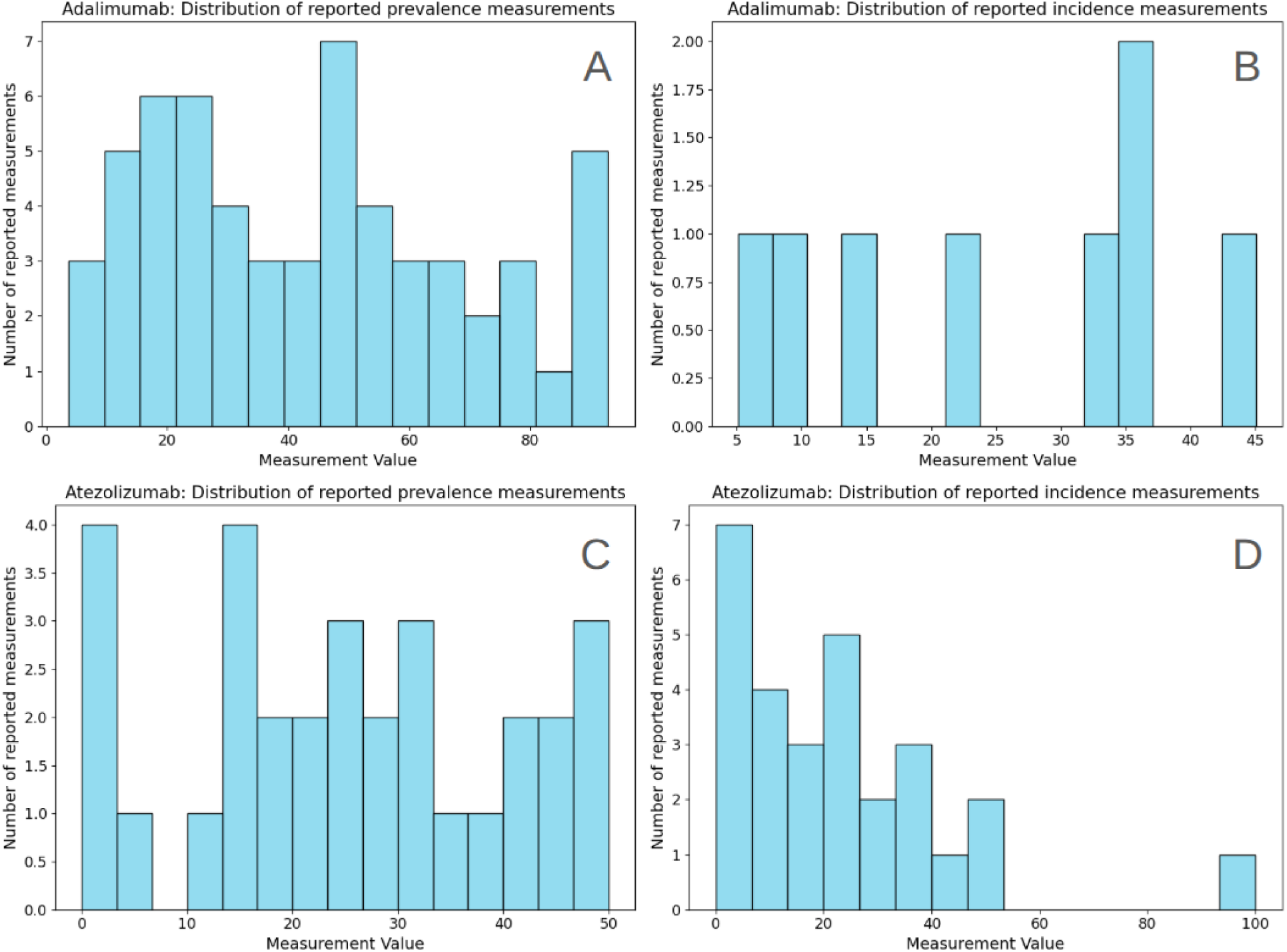
Immunogenicity variability of therapeutics. Variability of Adalimumab prevalence (A) and incidence (B). Variability of Atezolizumab prevalence (C) and incidence (D).

**Figure 9.**
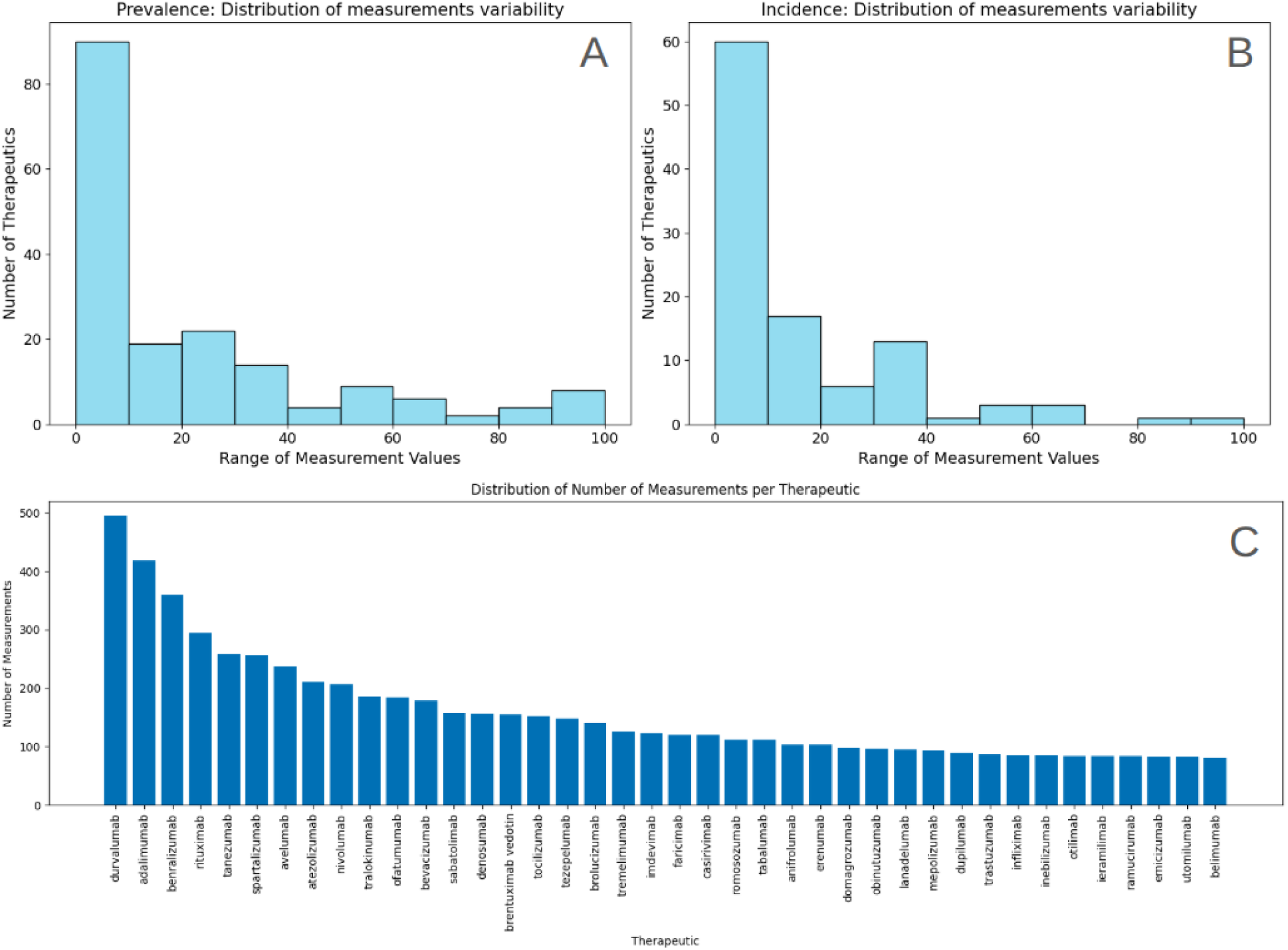
Immunogenicity variability of therapeutics. Distribution of variability defined as the difference between the maximum and minimum reported immunogenicity values for prevalence (A) and incidence (B). The number of raw measurements for top 40 most covered therapeutics (C) ordered by the number of reported measurements.

There might be several factors that influence the variability of the observed values. One ingredient is a dose dependent relationship with the observed ADA. Larger doses can be intuitively associated with higher exposure to the immune system, leading to larger observed ADA response. The relationship is non-trivial - for example in NCT01107457 - here the inverse relationship can be observed for Ixekizumab. Another notable example of a high variability report is the clinical trial with id NCT01307267 - where ADA against Utomilumab were measured. The results greatly varied across different cohorts, where different dosing regimens were implemented, but we have not observed any clear relationship to the dose. We believe the significant heterogeneity observed across cohorts in this study is related to small cohort sizes, further reinforcing the necessity of multiple studies to establish the robustness of the measurements. This is particularly crucial given the limited sample sizes inherent in Phase 1 trials, where conclusions drawn from insufficient data carry a high degree of uncertainty. Another aspect influencing the volatility of the measurements is the patient’s history. For example, in NCT03264066 there are clear and inconsistent differences between cohorts naive and exposed to treatments. The view is further convoluted by conflicting impact of various treatment context features. For instance, we noted a conflicting impact of comedications (Figure 10). Comedications with inhibitory effect on the immune system are expected to lower the ADA, but interestingly, we observed cases with inverse relationship (e.g. Avelumab in Figure 10).

**Figure 10.**
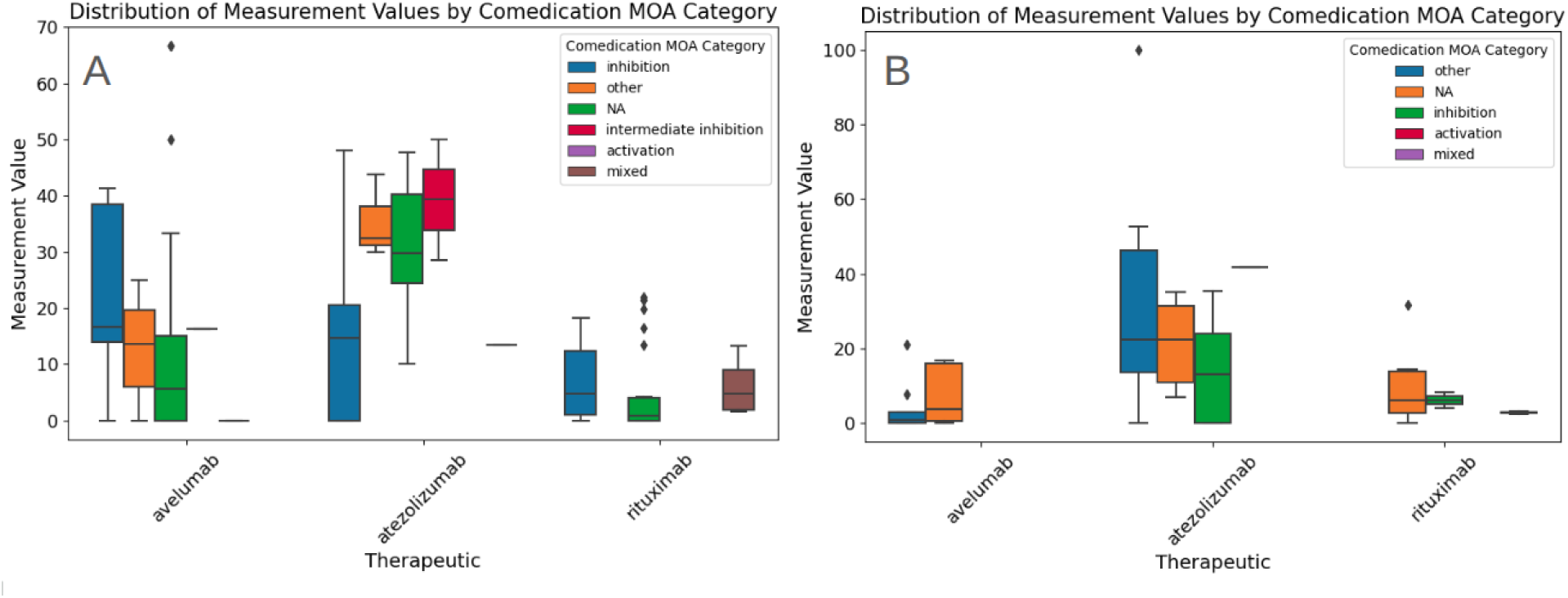
Comedication mode of action inconsistent impact on prevalence (A) and incidence (B) for selection of therapeutics.

Altogether, a wide range of measurements for a single molecule is a clear indicator that immunogenicity predictions from ‘sequence alone’ cannot yield good results. Clinical context appears to hold the key (Hu, Wu, and Swanson 2025) but as shown by our random forest exercise, the solution is not to simply aggregate everything into a simple model.

### Beyond the noise: the limited impact of therapeutic context

We established that sequence features alone or aggregating all the results in a simple machine learning algorithm do not yield generalizable results Therefore, we performed an analysis of the therapeutic context to explore the factors that might affect immunogenicity beyond molecular features alone and thus show a path towards better predictors.

Patient condition and route of administration are widely recognized as important variables influencing the immunogenicity of therapeutic proteins (Harris and Cohen 2024). The inflammatory environment might provide costimulatory signals required for T-cell activation. Route of administration may further shape immunogenic outcomes by affecting antigen processing and presentation: subcutaneous or intramuscular delivery could facilitate enhanced uptake by local antigen-presenting cells, while intravenous infusion may bypass some of these mechanisms. Consequently, understanding and optimizing the interplay between dose magnitude and route of administration may prove instrumental in mitigating ADA responses and improving overall clinical outcomes.

We assigned diseases reported for cohorts to a set of disease categories to explore the relationship between the patient state and ADA. We calculated the number of unique therapeutics and the distribution of measurements for each disease category (Figure 11). We noted higher ADA for autoimmune patients than in cancer patients. We repeated the same analysis for the reported routes of administration (Figure 12).

**Figure 11.**
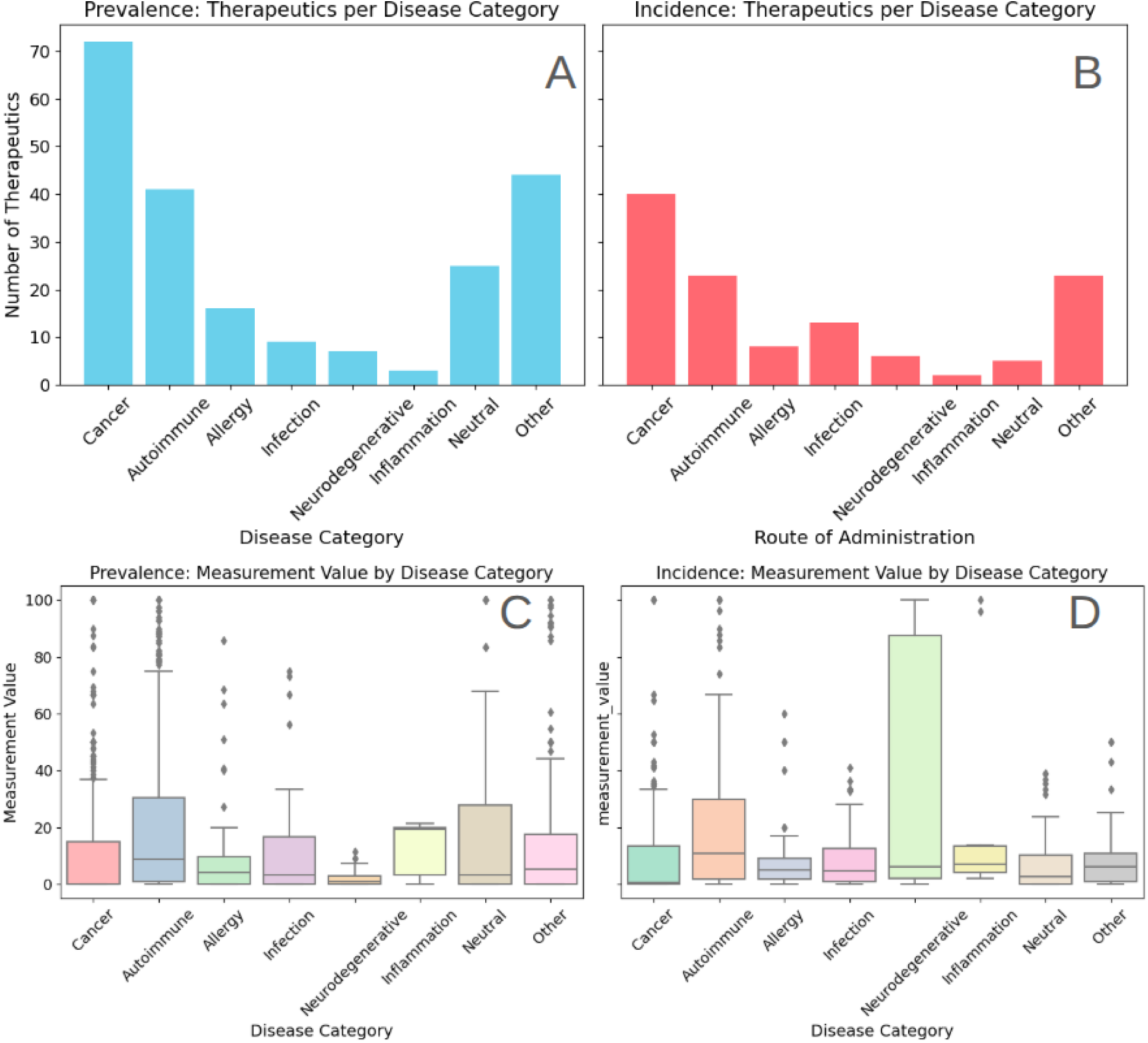
Number of distinct therapeutics used in various disease treatments for which we have collected immunogenicity measurements: Prevalence (A) and incidence (B); Distribution of immunogenicity prevalence (C) and incidence (D) for different categories of diseases.

**Figure 12.**
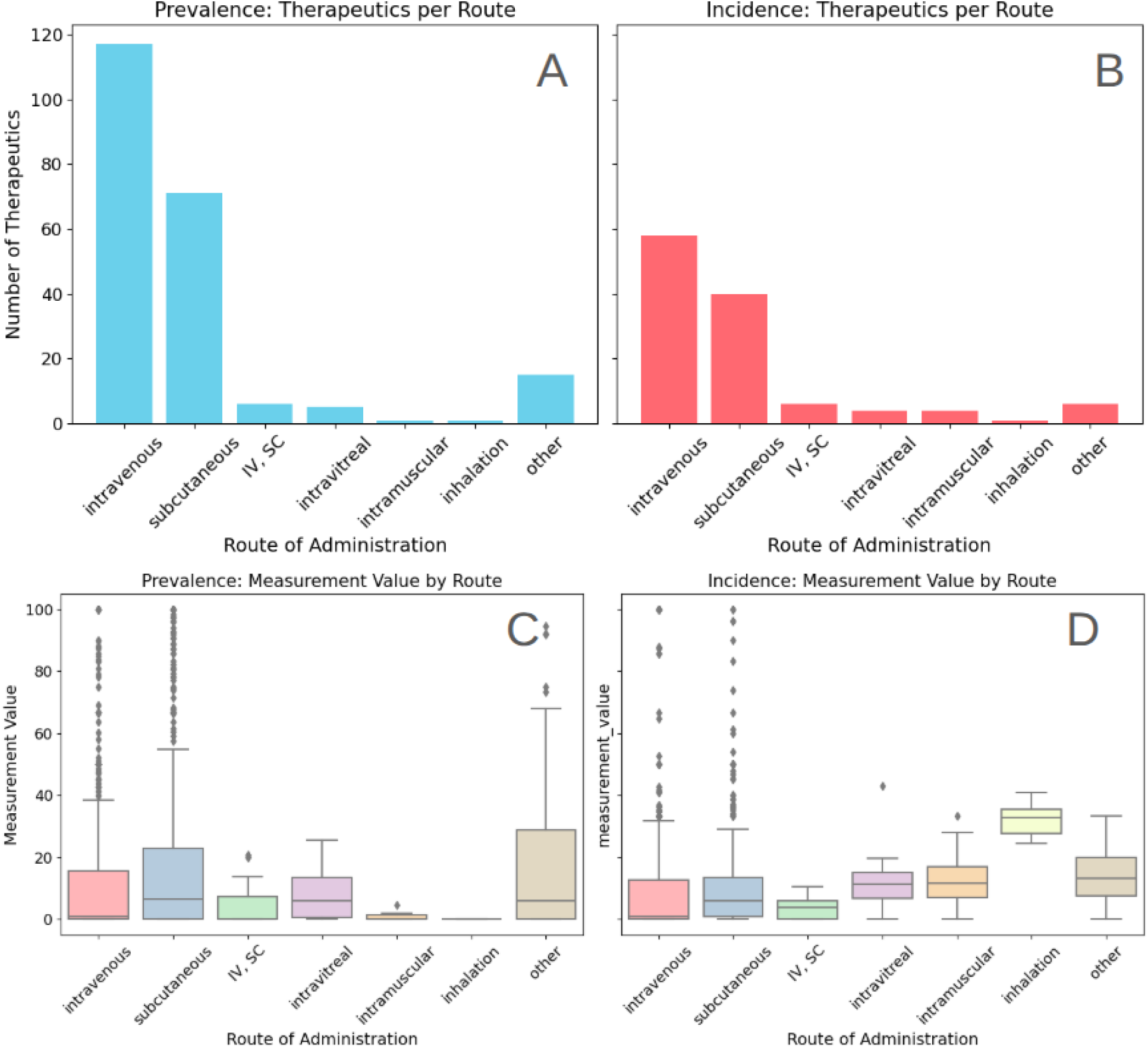
Route of administration: number of distinct therapeutics for various routes of administration in prevalence (A) and incidence (B); Distribution of immunogenicity in different routes of administration for prevalence (C) and incidence (D).

The analysis yielded two observations: reported route of administration is biased towards subcutaneous and intravenous routes with the latter consistently showing lower ADA in prevalence and incidence. Interestingly, we noted high ADA incidence in patients where the drug was administered by the inhalation but the amount of data in the category prohibits us from drawing any conclusions.

#### Therapeutic mode of action shows inconsistent impact on immunogenicity

Based on the results of (Hu, Wu, and Swanson 2025), we explored the impact of drug modes of action (MOA) on the ADA prevalence and incidence (Figure 13, Figure 14). The modes of action were assigned based on the interactions of therapeutics with the targets, as described in the methods section. We assigned the “activation” category when the therapeutic had direct stimulatory impact on the immune cells (e.g. CTLA4 antagonist). Similarly, the “inhibition” category was assigned when the interaction with the therapeutic target caused direct immune cells inhibition (e.g. B cell depletion agents: CD19-binding antibodies, CD20-binding antibodies; Co-stimulatory receptor blockers: OX40L (TNSF4) blocker). Surprisingly we have not observed any vertical separation between distinct modes of action categories.

**Figure 13.**
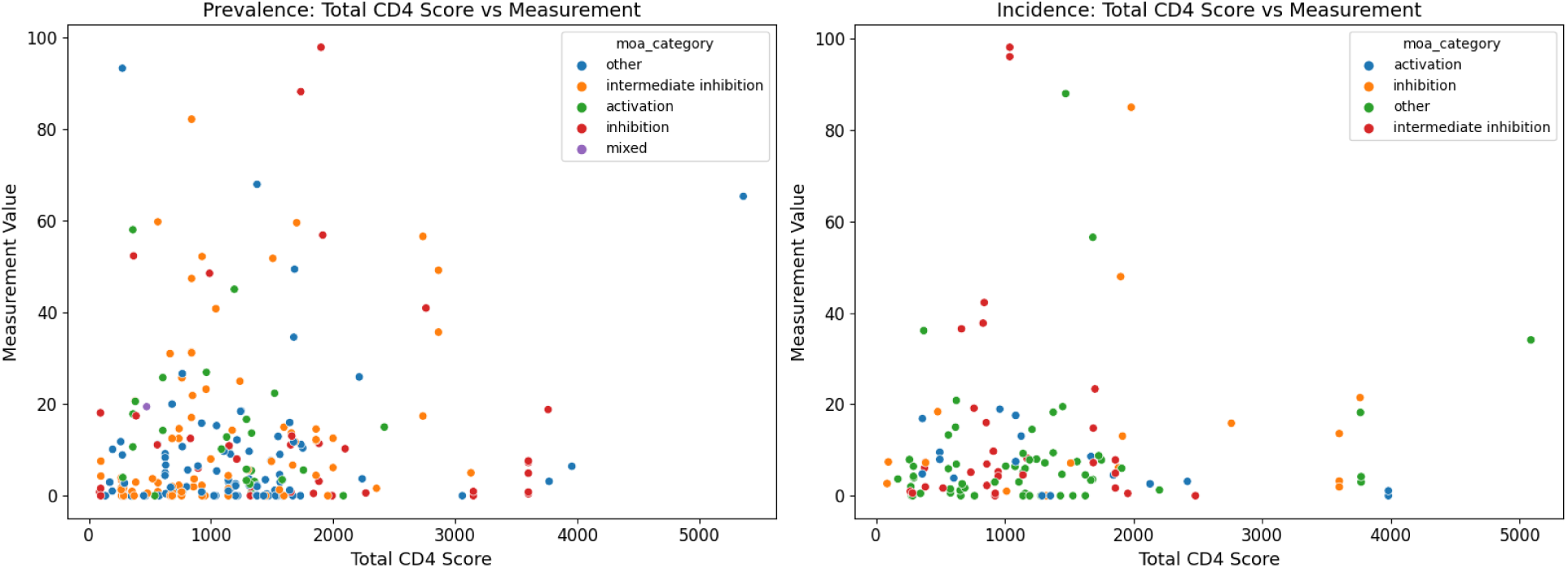
CD4 Immunogenicity score against ADA prevalence (left) and incidence (right) colored by the therapeutic mode of action.

**Figure 14.**
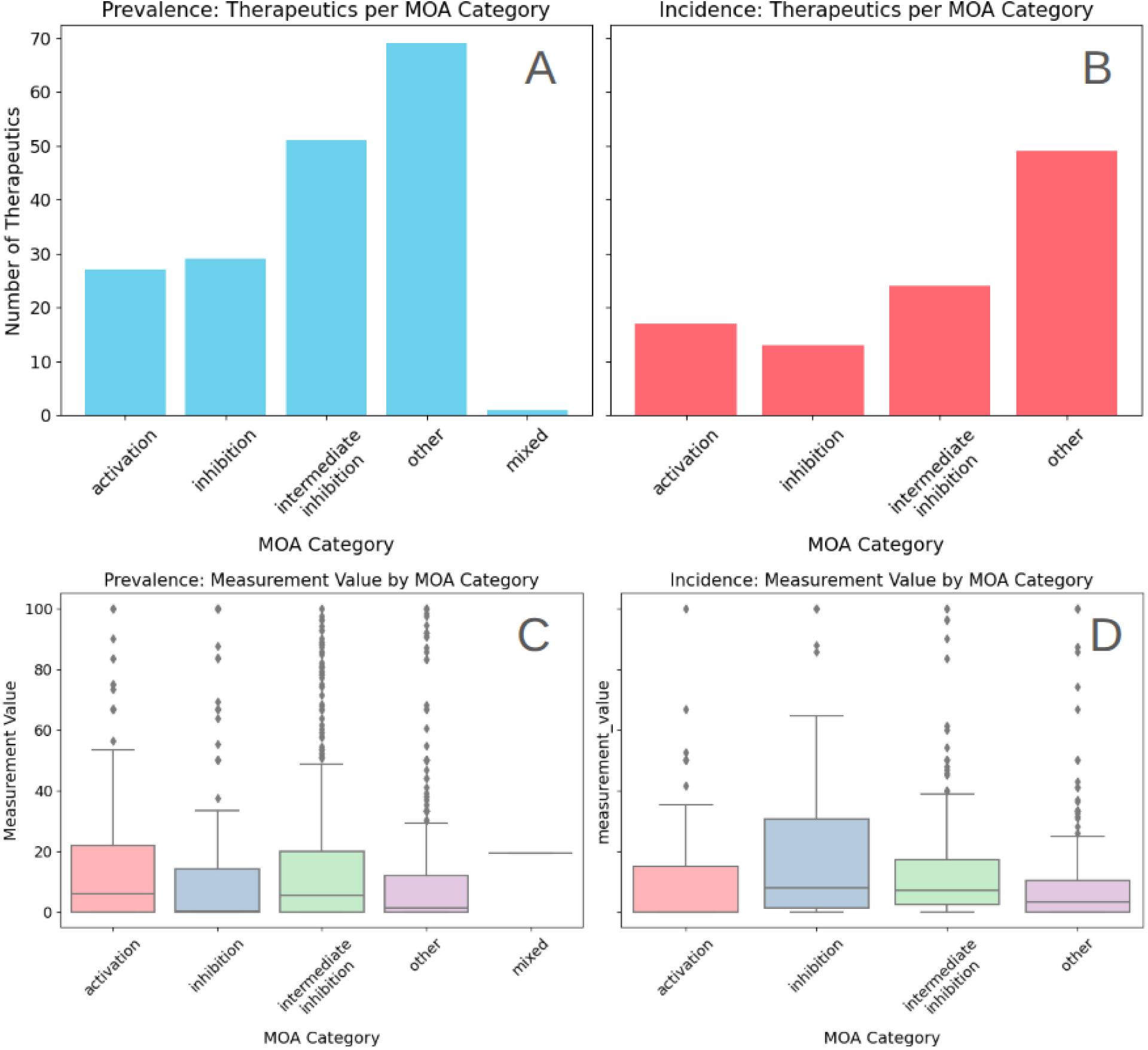
Therapeutic MOA: number of distinct therapeutics for each MOA category in prevalence (A) and incidence (B); Distribution of immunogenicity for various MOA in prevalence (C) and incidence (D)

Surprisingly, the observed ADA is not consequently associated with immune system activation or inhibition and is different for prevalence and incidence. The finding does not rule out the results from (Hu, Wu, and Swanson 2025) - it merely shows that it is a much more complicated problem.

#### Concomitant immune system inhibitors moderately reduces the risk of immunogenicity

To further explore the patients state impact on ADA, we annotated non antibody interventions for each cohort. We hypothesized that those co-medications can act as B or T cells activators or inhibitors can cause the differences in the ADA incidence and prevalence. The results are shown on Figure 15.

**Figure 15.**
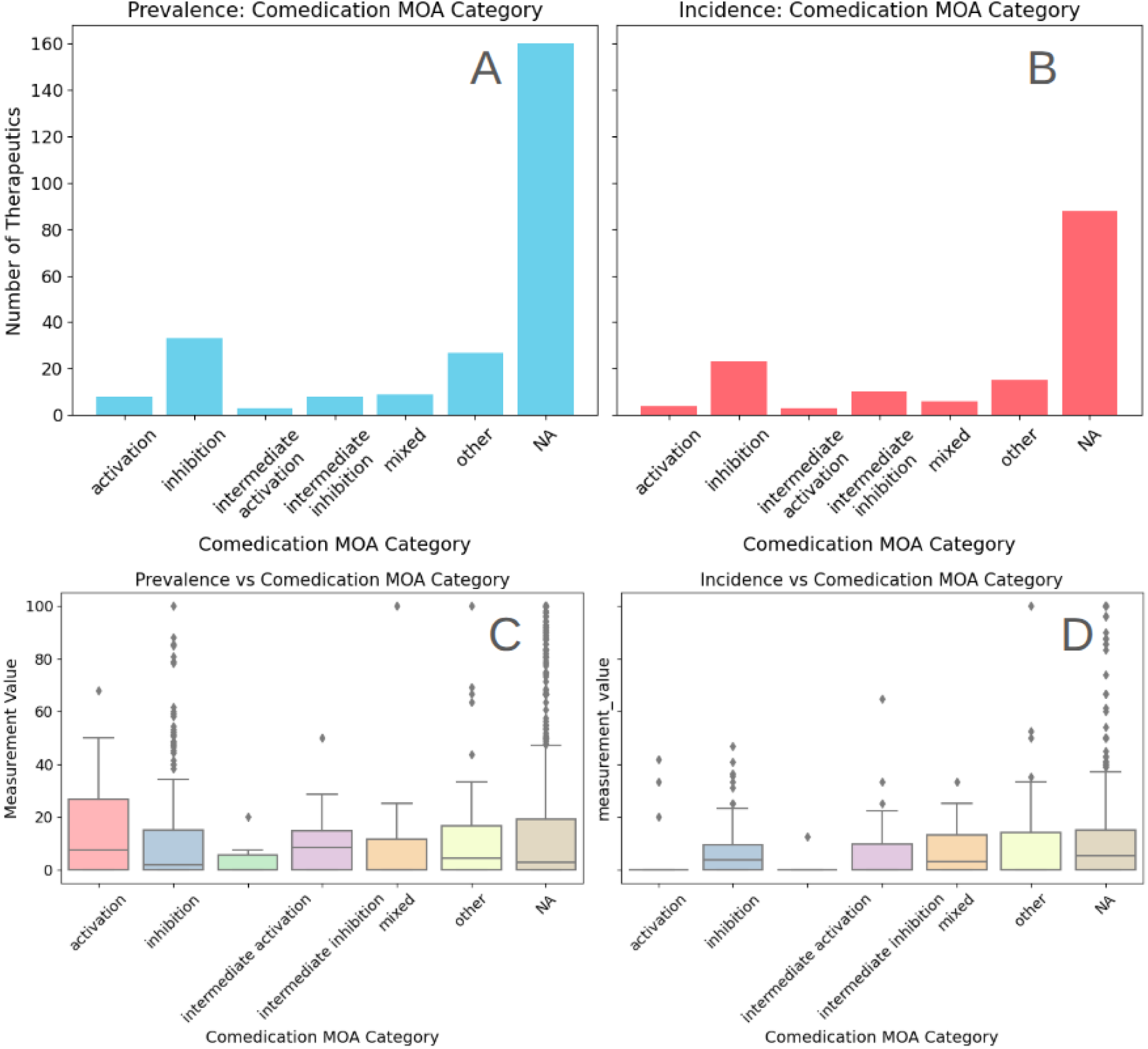
Comedications MOA: Number of therapeutics for which we registered additional interventions in prevalence (A) and incidence (B). Immunogenicity in groups with different comedications MOA for prevalence (C) and incidence (D). Comedication immune system inhibition seems to reduce immunogenicity of the therapeutic in prevalence but not for incidence, which we attribute to a small number of data points.

The MOA assignment of the co-administered non-antibody interventions is described in the methods. We noted that the mean immunogenicity is lower for prevalence in treatments where immune system inhibitors are introduced as a concomitant intervention, compared to treatments with activating comedications..

#### Presence of the Fc region is associated with reduced immunogenicity

Each therapeutic in the database has been assigned to one of two structural categories: Fc containing antibodies (formats with Fc and antigen-binding fragments) and antibody fragments (antigen-binding, no Fc region, e.g., scFv, Fab). As we explored the immunogenicity in structural formats, we noted that the majority of existing measurements corresponds to the antibodies with a classic format - consisting of two heavy and two light chains (Figure 16). While whole classical mAbs also meet the criteria of Fc containing antibodies, we assigned them a distinct category, separated to study engineered formats’ safety advantages over natural ones.

**Figure 16.**
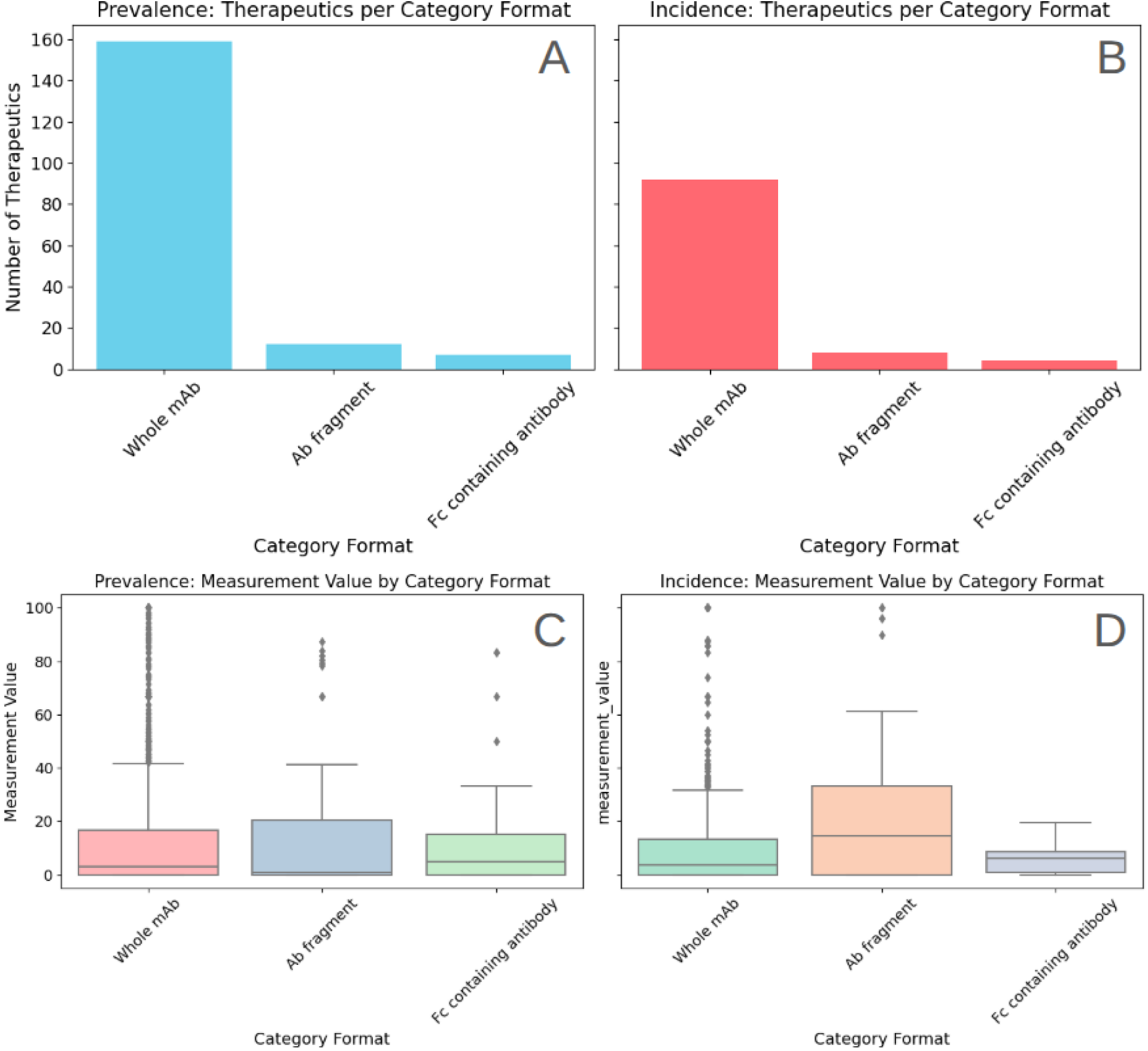
Therapeutic format: number of distinct therapeutics for various formats for prevalence (A) and incidence (B) measurements; Immunogenicity prevalence (C) and incidence (D) for different structural categories.

The Fc region of a therapeutic antibody influence on immunogenicity stems primarily from its interaction with the host’s immune system via Fc receptors and the complement system. Interestingly we do not observe a notable difference in immunogenicity of engineered formats containing Fc. While the average immunogenicity incidence of antibody fragments is noticeably higher than in Fc containing antibodies (whole mAbs and other formats), the coverage of novel formats is too sparse to draw any meaningful conclusions.

Since there are multiple Fc receptors which mediate the interaction between the therapeutic and the patient’s organism, we visualized the immunogenicity for different C germline genes assigned by RIOT (Figure 17). The therapeutics are heavily biased towards IGHG1 and IGKC. While less abundant, IGLC2 shows lower mean immunogenicity for both prevalence and incidence. We attribute unusually IGHG2 high ADA prevalence to the complement activation, although the results here also should be interpreted with caution because of the small number of examples.

**Figure 17.**
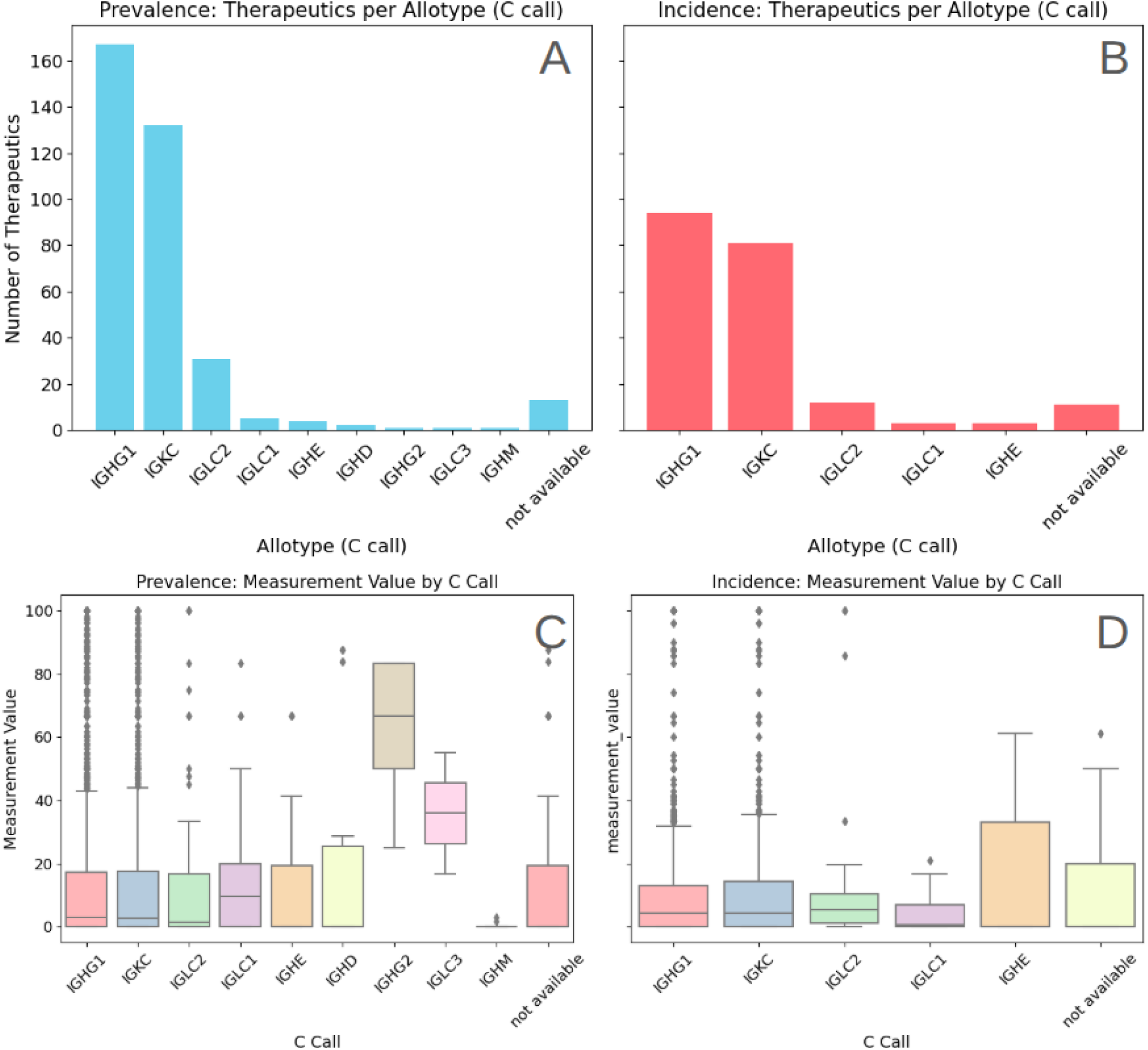
C genes: number of distinct therapeutics for C genes assigned for prevalence (A) and incidence (B) measurements; Immunogenicity prevalence (C) and incidence (D) for different C genes categories.

## Discussion

Currently, the gold standard for preclinical immunogenicity assessment is MAPP (MHC-associated peptide proteomics) (Karle 2020) and T-cell activation assays (Cohen and Chung 2021). These are in vitro analyses in which proteins are cleaved and presented on MHC II in the presence of T cells, examining either which peptides bind to the MHC complex or whether T cells are activated. This approach is time-consuming, hence the need for in silico methods that could speed up the process. At present, advanced computational methods focus on specific stages of ADA formation: predicting peptide binding to MHC (Reynisson et al. 2020) and predicting T-cell immunogenicity (Wang et al. 2023; Dhanda et al. 2018), i.e., the ability to activate T cells. Models for predicting antigen processing concentrate on MHC I, and results for MHC II lag far behind (Farriol-Duran et al. 2023). Antigen processing is central to immune responses, yet is profoundly influenced by factors such as the type and maturation state of antigen-presenting cells and the surrounding environment, making it challenging to predict which specific epitopes will ultimately be displayed on MHC II molecules.

The computational methods that would evaluate the “overall” immunogenicity of therapeutic proteins under clinical conditions are not well developed: as we showed in this study, humanization provides only limited results and is not backed by a mechanistic model of our understanding of the problem. Meanwhile, models that assess immunogenicity do not fully explain the sources of its emergence. For instance, in study (Hu, Wu, and Swanson 2025), the authors created a binary classifier to predict the occurrence of ADA (whether or not the percentage of participants with ADA were greater or equal 10%). Analysis of the results indicated that apart from the presence of the T-cell epitopes, the mechanism of action (T-cell activation/suppression) was a key factor in immunogenicity. With the database of much bigger coverage, we were not able to detect similar patterns.

There have been studies investigating the immunogenicity of T-cell epitopes and prediction of T-cell responses. The majority of the research, however, focuses on neoantigens in the context of vaccines, where immunogenicity is a desired property. While many analogies can be drawn, this is evidently a distinct case. In specific, using tools such as NETMHCPan (Reynisson et al. 2020) is misunderstood. Considering predicting the immunogenicity of the vaccines, it is sufficient to find any matching MHC in patients, therefore looking only at the popular alleles which covers 90% of the population is sufficient. When considering the immunogenicity of the therapeutic proteins, however, all alleles should be examined (or at least the abundant in the target population).

As we concluded from the analysis, sole exposition of immunogenic peptide is not sufficient to trigger immune response, and other factors need to be considered, beyond biophysical properties analyzed in this work. Notably the antigen processing, T-cell activation signals and the immune tolerance mechanisms. T-cells need two activating signals: from TCR and costimulatory from CD28 activation. Lack of co-stimulation renders T-cell anergic, as represented by lower TCR expression and inhibited proliferation in response to antigen. Yet If T-cell gets activated then, preserving tolerance is ceded to regulatory lymphocytes, which reduce T cell costimulatory signals by binding to CD80 and CD86 on antigen-presenting cells, and they further downregulate immune responses through the secretion of anti-inflammatory cytokines such as IL-10 and TGF-β.

To further complicate the image, the presence of Tregitopes (Anne S. De Groot et al. 2008) should be evaluated. Tregitopes are primarily derived from the Fc region of IgG but also appear in conserved framework regions, suggesting they may broadly contribute to tolerance by inducing the expansion and activation of Tregs when presented on MHC class II molecules. From a therapeutic standpoint, regulatory T-cells (Tregs) in theory can be harnessed to induce tolerance in autoimmune diseases. In such conditions, Tregs can be expanded or induced (e.g., through FOXP3 expression) to dampen pathogenic immune responses (Gołąb, Lasek, and Jakóbisiak 2017). Tregs can also perpetuate tolerance through a process called “infectious tolerance”, wherein they secrete IL-35 to convert other T cells into Tregs. However, in the context of cancer, these same mechanisms can be detrimental, because Tregs may contribute to tolerance of tumor antigens, effectively protecting cancer cells from immune attack (Gołąb, Lasek, and Jakóbisiak 2017). Therefore, while Tregs and Tregitopes could be beneficial in treating autoimmune diseases, they must be carefully managed or blocked in cancer therapy.

The presence of a “warning” signal, which is mediated by PAMP and DAMP can trigger an immune reaction which underscores the importance of patient conditions as an important factor in predicting the risk of immunogenicity. Furthermore viral infections might alter how antigen is processed, therefore uncover hidden T-cell epitopes.

Our analysis reveals that the reported immunogenicity of therapeutics is highly variable, a complexity driven by several interconnected factors. This variability cannot be attributed to a single cause. We found that the relationship between drug dosage and anti-drug antibody (ADA) response is often unpredictable, in cases of immune system inhibition, showing an inverse correlation where one might expect a direct one, as larger doses increases therapeutic exposure to the T-cells. The picture is further complicated by patient-specific factors. For example, a patient’s prior treatment history (treatment-naïve vs. exposed) can lead to inconsistent ADA responses between cohorts. Similarly, the impact of comedications can be paradoxical; immunosuppressants, which are expected to lower ADA rates, were occasionally associated with an increase. Crucially, this inherent volatility is amplified by the small cohort sizes common in early-phase trials. Small sample numbers make it difficult to distinguish true biological effects from statistical noise, leading to uncertain and often unreliable conclusions from a single study. Those factors strongly underscore that a comprehensive and reliable understanding of a therapeutic’s immunogenic profile can only be achieved by aggregating and analyzing data from multiple, diverse clinical trials.

Finally, there are multiple factors that impact the quality of data. There may be substantial differences in how studies measure immunogenicity in terms of protocols and assay sensitivity. Product-related factors that are not captured in the database, such as impurities or hydrophobicity, might lead to aggregation further complicating the comparisons.

The pursuit of reliable in silico immunogenicity prediction is slowed down by a reductionist focus that ignores crucial layers of biological complexity. The current paradigm, centered on a simplified T-cell epitope prediction pipeline, is built on a foundation of volatile clinical data and is weakened by the unresolved problem of B-cell epitope prediction. Each step of its core cascade - from modeling proteolytic cleavage to assuming activation from MHC binding - is loaded with inaccuracies. Most critically, this approach fails to account for the powerful modulatory effects of the patient’s unique context, including their genetic makeup (HLA and FcR polymorphisms), the state of their immunological tolerance, and the specific inflammatory environment of their underlying disease. The clinical reality that a single therapeutic can have vastly different immunogenicity profiles in different diseases is strong evidence that a protein-centric model is insufficient.

Making meaningful progress requires a fundamental paradigm shift away from this linear, component-based methodology towards an integrated systems immunology framework. Future predictive models must evolve to become multi-parameter systems that take as input not only the features of the therapeutic molecule but also patient-specific variables. Such a model would integrate data on the protein’s sequence and potential epitopes with the patient’s HLA and FcR genotypes, their disease state (and its associated inflammatory profile and co-medications), and their baseline tolerance status. By addressing the complexity of the host-drug interaction, we can move from basic risk categorization towards the (accurate) personalized immunogenicity prediction.

## Methods

### Data format

The data is formatted as .csv into several sheets, akin to Excel. This was dictated by the fact that a single therapeutic needs to be associated with multiple values in a relational database fashion, which is not suitable for a single .csv file. Sharing dumps of relational databases though is not user friendly. The entity relationships diagram of the database tables is presented on Figure 18.

**Figure 18.**
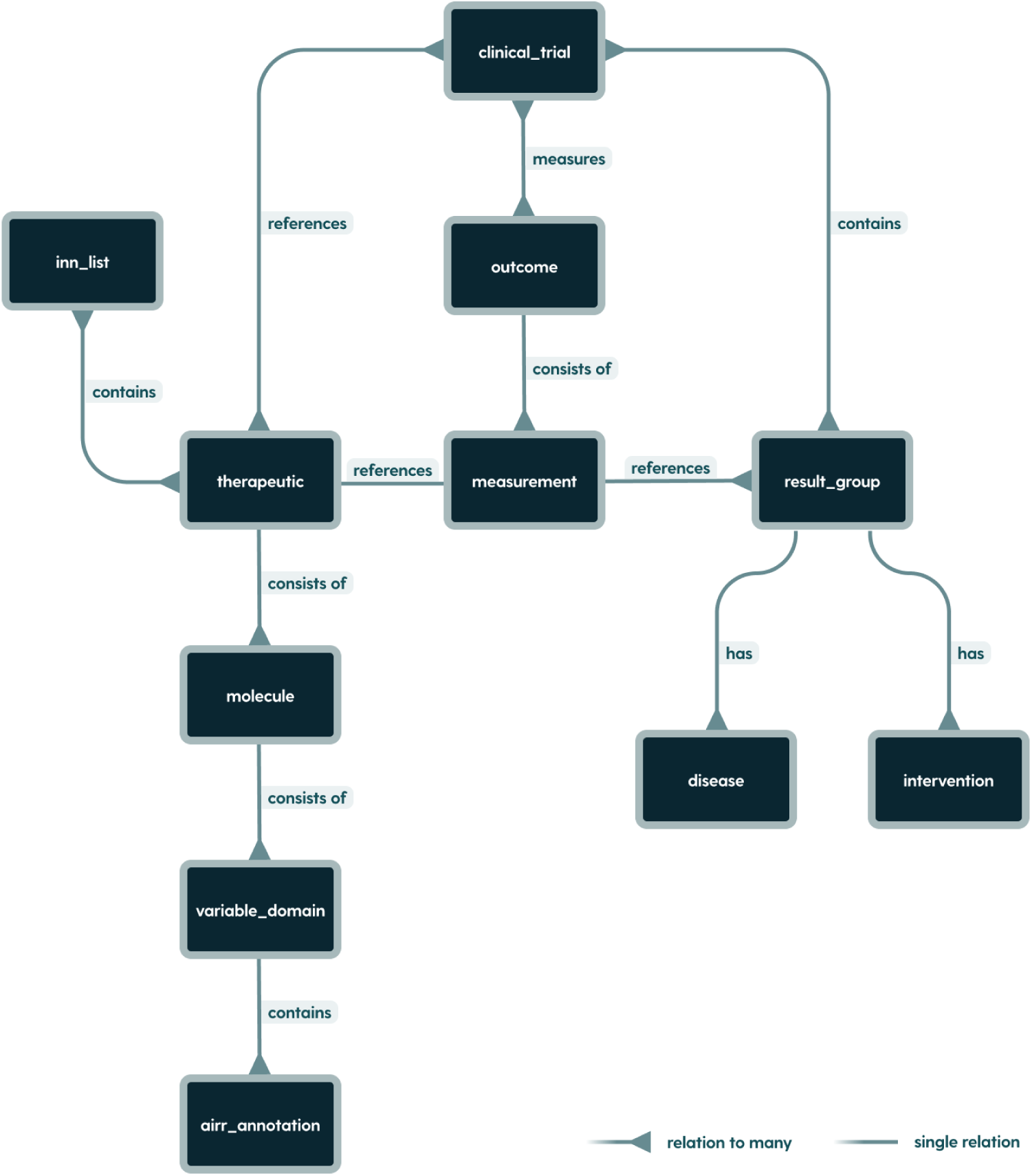
Entity relationship diagram of the database tables.

At the core of the database is a therapeutic table, which contains drug name, its synonyms, information about its format and post-translational modifications. Each therapeutic is sourced from International Nonproprietary Names (INN) Lists, so references to those are stored in the inn_lists. Each therapeutic consists of one or more amino acid chains, which are stored in the molecule table, as full sequences, together with their target antigens. Each molecule may contain variable domains, which are stored in the variable_domain table, with the position of the domain in the sequence. For example a typical monoclonal antibody, which is built from heavy and light chains, will have 1 record in the therapeutic table, two records in the molecules table and two records in the variable_domain table (one assigned to each molecule). On the other hand, an scFv therapeutic will have only one molecule with multiple variable regions assigned to it.

Each therapeutic may have some clinical trials assigned to it in which it was tested. The table clinical_trial contains basic trials information, such as title and phase. As the relationship between therapeutic table and clinical_trial table is many to many, it is realized with a third table holding references from both tables (therapeutic name and clinical trial identifier).

Each clinical trial measures various outcomes. We focus on the outcomes investigating immunogenicity. The immunogenicity may be measured at multiple points in time, so each distinct point in time should have a separate record in the outcomes table. The immunogenicity in clinical trials is measured for each cohort separately. Each cohort will have associated diseases and interventions in the corresponding tables. Interventions table contains information about the treatment regimens if they are present, including the dosage and the route of administration. Finally, the measurement table contains immunogenicity measurements associated with corresponding clinical trials outcomes and cohorts. The values were properly annotated to facilitate inter-study comparisons. For a more detailed description, see the following chapter. From all of the above tables we derive three datasets presenting incidence, prevalence and pre-existing immunogenicity.

### Curation of the database

We manually review INN Lists, published by WHO to identify antibody therapeutics and extract full sequences (including constant regions), target information and post-translational modifications. Molecules are numbered via the IMGT scheme using RIOT, with variable domains described in the AIRR Format (AIRR Community, n.d.). Variable regions are manually grouped together by the Fv, where applicable, so that in cases of multiple targets it is possible to assign the target to each Fv separately. Trade names are imported from OpenFDA <u>(OpenFDA)</u>, synonyms from the NCI Thesaurus <u>(National Cancer Institute (.gov) 2011)</u>.

A database of antibody immunogenicity measurements was constructed by compiling monoclonal antibody names and synonyms, then importing relevant clinical trials from ClinicalTrials.gov into tables such as clinical_trial, outcome, result_group, and measurement. Clinical trials report multiple outcomes (toxicity, pharmacodynamics, immunogenicity) at specific time points per cohort (result_group). We focus only on immunogenicity related outcomes.

Cohorts were annotated with diseases and interventions, so that each cohort has assigned diseases and treatment interventions, together with dosages and routes of administration (if present). Diseases were further categorized and mode of action was assigned to the interventions, for both: antibodies and comedications.

Given the heterogeneity in how studies reported their results, several strategies were implemented to standardize the data. When multiple measurements for a single outcome were reported at different time points, artificial outcomes were created so that each outcome corresponded to a single time point. Neutralizing antibody responses were specifically flagged, allowing for their separation from other immunogenicity measurements. In instances where partial results were reported, a grouping key was assigned to facilitate proper aggregation. An example of partial measurements are measurements of immunogenicity at different titer levels or performing more accurate measurements after the screening.

Each measurement was annotated with a type indicator to distinguish between positive, negative, missing or inconclusive outcomes. Additionally, studies that reported overall immunogenicity alongside relative measures, such as treatment-emergent or treatment-boosted responses, were annotated accordingly. This approach enabled direct comparisons, with immunogenicity prevalence defined as the sum of pre-existing, treatment-boosted and treatment-emergent responses.

Further data harmonization included adjusting measurement group sizes where the number of eligible patients differed from the initially defined cohort size. In some cases eligible patients were extracted from metadata; in other cases they were extracted from the measurement description; in the remaining cases eligible patients were derived by subtracting missing and inconclusive participants from the original cohort size. This unification allowed for another unit conversion - measurement units were normalized to represent the percentage of participants, ensuring consistency across studies. In cases where studies differentiated between transient and persistent measurements, transient responses (defined as positive at one time point and negative at another) were flagged and annotated with a grouping key to allow for correct summation with overall ADA scores.

With the above annotations we constructed three additional derived datasets: presenting immunogenicity incidence, prevalence and baseline for cohorts in which only one antibody therapeutic was administered. For all three datasets, after we grouped partial measurements and unified the units (participants were converted to percentage values based on identified groups sizes), we grouped the measurements from multiple points in time, so that each cohort had a single immunogenicity value, assigned disease and intervention. The results were further weighted averaged to produce one measurement for each therapeutic, disease and route of administration. For the downstream analyses, only the measurements from studies completed in 2016 or after were selected.

#### Therapeutic format

In our database we proposed a unified system for INN present antibody based drugs. We introduce a structural format which focuses on the structure of the antibody therapeutic. We distinguish 115 distinct structural formats with the examples of: immunoglobulin G1 scFv-h-CH2-CH3(_scFv)_h-CH2-CH3 or immunoglobulin G1-kappa with crossed domains only name a few. Then we classified each therapeutic to the functional format, focusing on the function where there are classical bispecific antibodies (antibodies that can bind to two distinct epitopes), in the case of our database “bispecific” is for example faricimab - antibody targeting two distinct antigens: VEGF-A and ANG-2. However, if antibody is targeting two different antigens, but binding to one or more of the antigens posses a several additional function of the antibody such as T-cell activation or NK cells activation we classified that to the one of the newer formats such as BITEs (Bispecific T-cell engagers), DARTs (Dual-Affinity Retargeting proteins) or TrIKEs (Trispecific killer cell engager). Overall, we assigned 54 distinct functional formats.

Finally, each therapeutic was classified into one of the following following category formats: whole mAb, antibody fragment, Fc-containing antibody. We define the whole mAb as the “classical” tetramer molecules, consisting of 2 heavy and 2 light chains. Antibody fragments are antigen binding segments of an antibody (e.g. scFV, Fab), that does not contain Fc region. Fc-containing antibodies are all formats that include Fc and antigen binding fragments. While whole mAbs are also Fc-containing antibodies we assigned them to a separate category to investigate whether or not engineered formats have advantage over natural in terms of safety.

#### Target and therapeutic mode of action

Recent work (Hu, Wu, and Swanson 2025) highlighted that lymphocyte activation is a primary driver of antidrug antibody formation, indicating that immune-stimulating therapies may pose a higher immunogenicity risk. To explore whether the therapeutic mode of action also influences a biologic’s immunogenic potential, we categorized each interaction between the therapeutic and the target proteins. Each therapeutic with its matched target was individually assessed for the mode of action (MOA) type and MOA category. Definitions of MOA types and categories are shown in Table 2 and 3, respectively.

**Table 1.** Popular therapeutic targets. Target assignment was performed based on the molecule recognized by the variable regions to encompass multispecifics. Only targets with 10 or more distinct antibodies addressing them are given.

| Target | Number of therapeutics |
| --- | --- |
| CD3E | 61 |
| PDCD1 | 54 |
| EGFR | 34 |
| ERBB2 | 28 |
| PDL1 | 24 |
| CTLA4 | 23 |
| Spike glycoprotein | 21 |
| MS4A1 | 19 |
| CD40 | 16 |
| IL17A | 14 |
| TNFRSF17 | 14 |
| VEGFA | 13 |
| ALB | 13 |
| LAG3 | 13 |
| TIGIT | 13 |
| APP | 12 |
| CD19 | 11 |
| TNFRSF9 | 11 |
| MET | 11 |
| ERBB3 | 11 |
| PCSK9 | 10 |
| IL13 | 10 |

**Table 2.**
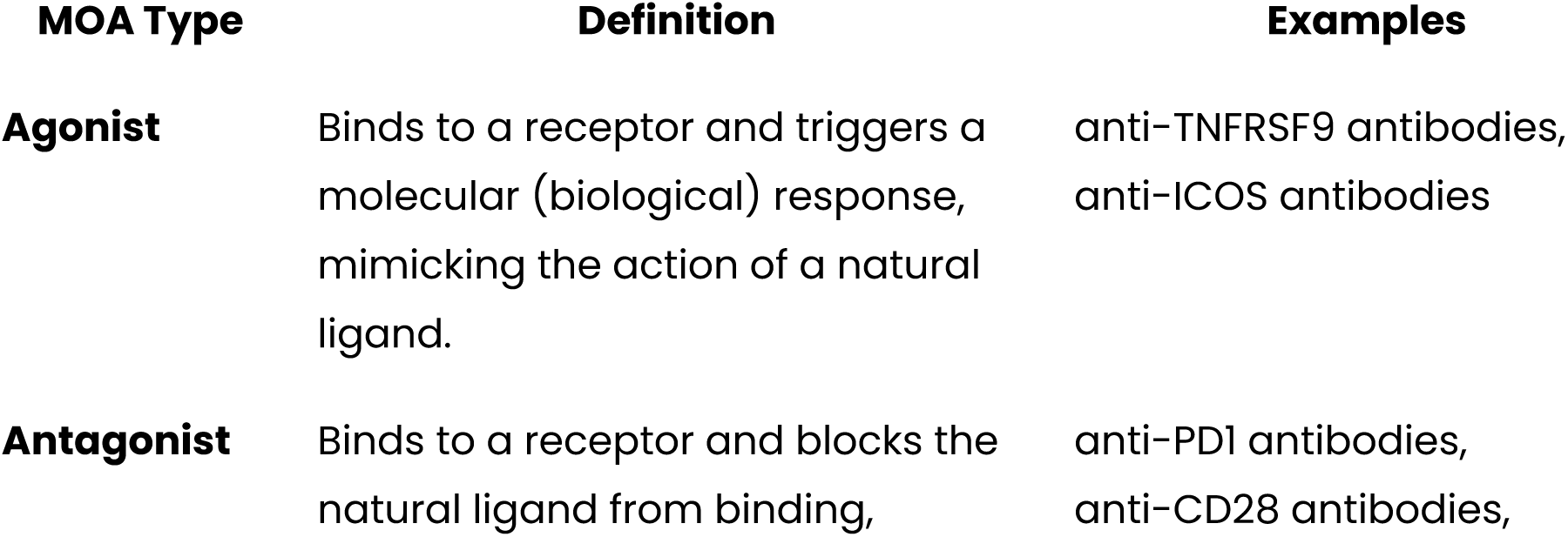
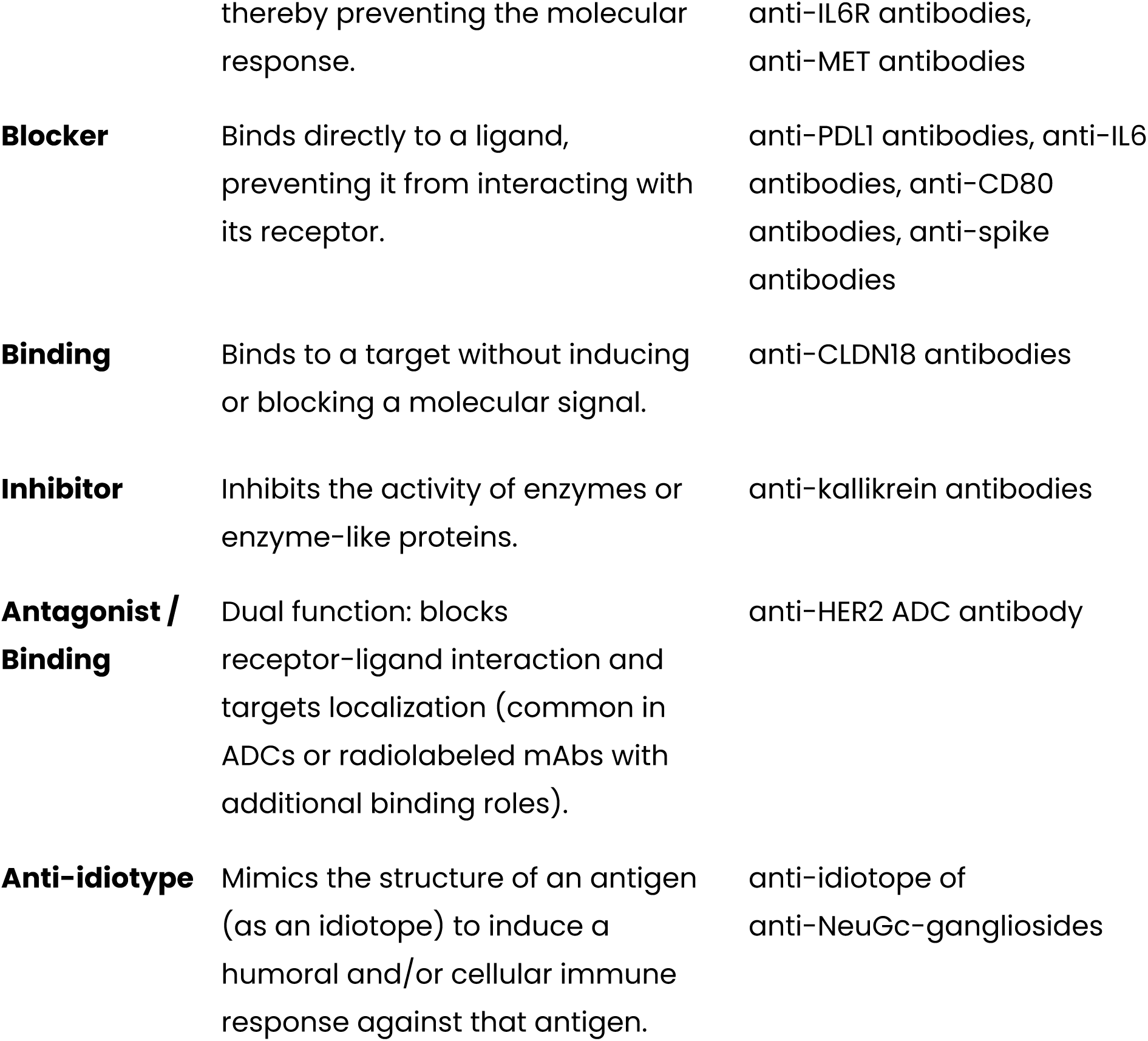
Target interaction types.

**Table 3.**
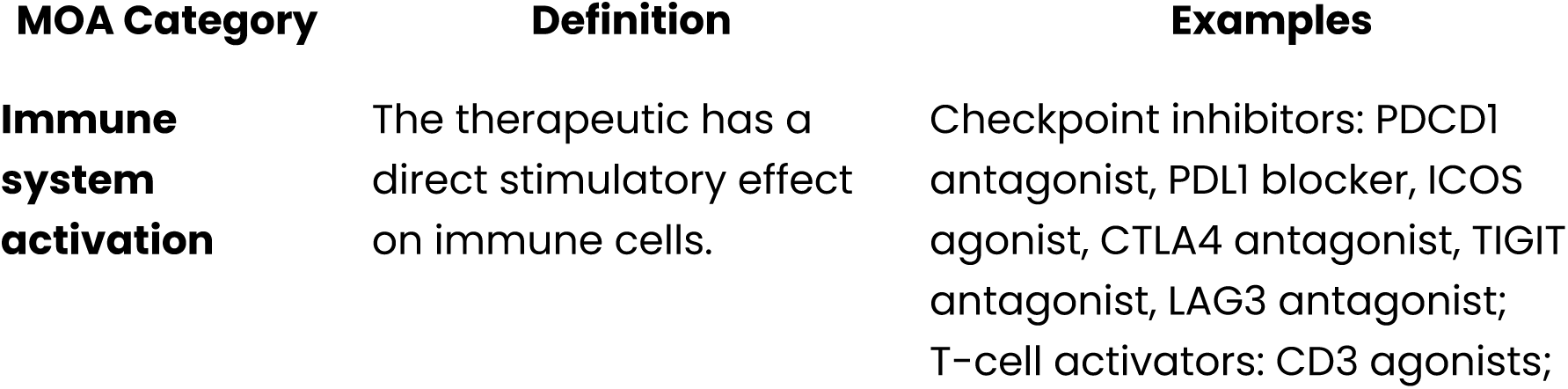
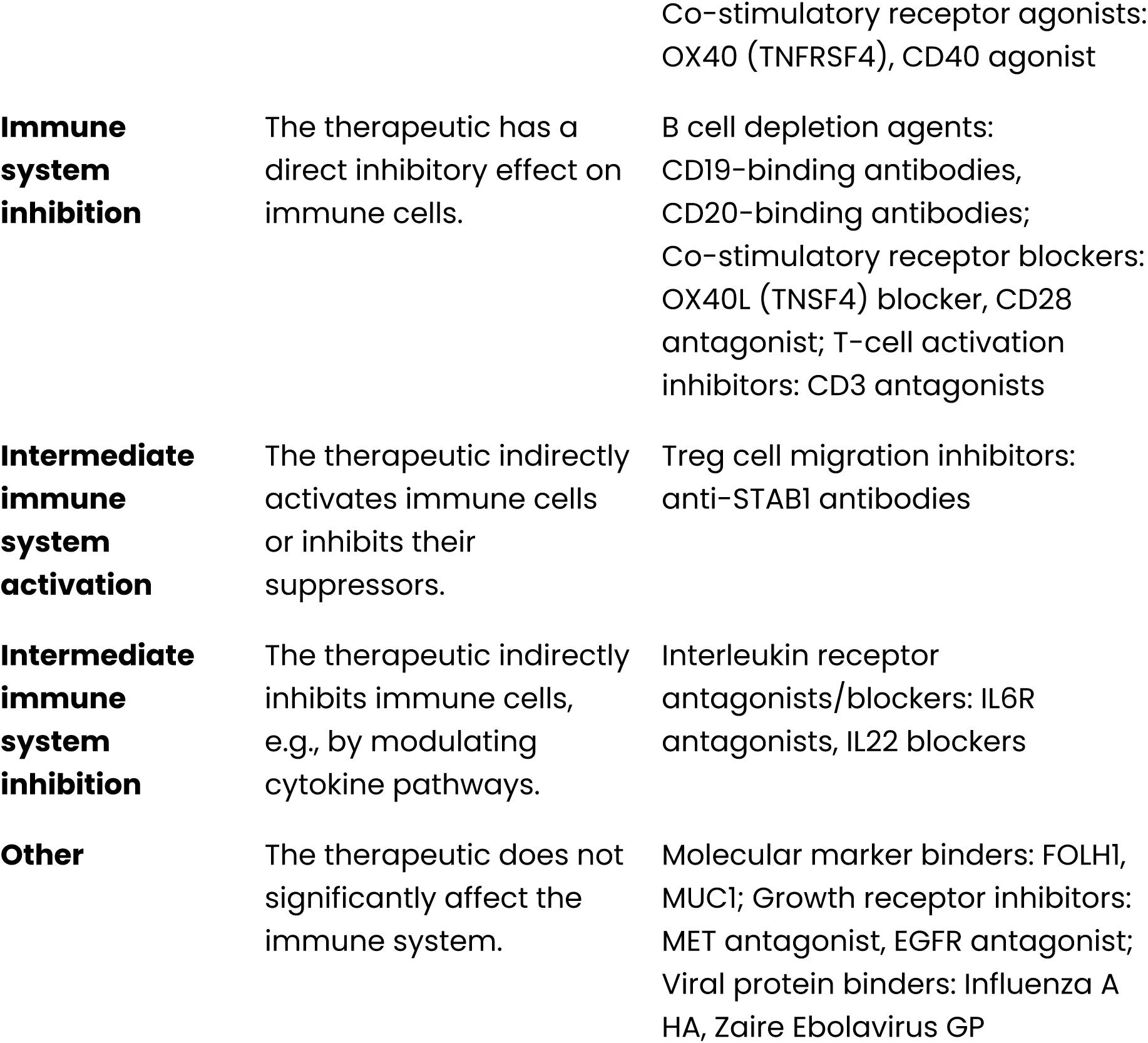
Modes of action inferred from therapeutic targets interactions.

### Co-medications mode of action

We classified non-antibody interventions used as comedications with antibodies in clinical trials. We hypothesized that comedications that can act as B-cells or T-cells activators or inhibitors can cause the differences in measured ADA incidence and prevalence. To address this, we prepared three annotations for co-medications: MOA, drug type and its main use. MOA of the co-administered non-antibody interventions is divided into seven main categories, described in the Table 4 below. Drug type is an assigned category describing the mechanism by which the drug causes its therapeutic effect. Main use is a one-word description of the clinical use of a given therapeutic. This is important for later analysis in the context of immunosuppressive drugs and chemotherapy with regard to their effect on ADA.

**Table 4.** Co-medications modes of action.

| **MOA Category** | **Definition / Mechanism** | **Examples** |
| --- | --- | --- |
| **Immune system activation** | Direct stimulation of immune cells. | IL-2 agonists, TLR agonists, cancer vaccines |
| **Immune system inhibition** | Direct inhibition or suppression of immune cells. | BTK inhibitors, calcineurin inhibitors, anti-CD20 antibodies |
| **Intermediate immune system activation** | Indirect activation of immune cells via modulation of cytokines or signaling molecules. | IL-2/IL-12 conjugates, TGF-beta inhibitors |
| **Intermediate immune system inhibition** | Indirect suppression of immune cells through signaling interference or environmental modulation. | Adenosine receptor antagonists, chemokine receptor antagonists, proteasome inhibitors |
| **Other** | No significant influence on the immune system; minor immunomodulatory effects not clinically relevant for classification. | NSAIDs, analgesics, antiemetics |
| **Many** (for regimens) | Multiple or mixed immunological effects, typically within combination chemotherapy regimens. | R-CHOP, FOLFOX |
| **NOT SPECIFIED** | MOA could not be determined due to vague labeling in clinical data (e.g., “supportive care” or “baseline therapy”). | Supportive care, baseline therapy |

### Identification of immunogenic T-cell epitopes

In order to evaluate how the presence of immunogenic epitopes shapes the image of clinical immunogenicity, we predicted T-cell immunogenicity scores for variable regions. The sequences were split into overlapping 15-mers and each segment was scored by the IEDB CD4+ immunogenicity prediction tool (Dhanda et al. 2018). The score for each amino acid position was calculated as a mean value of scores covering the residue. Sequence scores were obtained by summing all of the residue-level outcomes. To account for the well tolerated segments, we zeroed the scores of peptides observed abundantly in sequencing experiments in human studies from PairedAbNGS (Pawel Dudzic et al. 2025). We noted that the majority of therapeutics (even ones without any immunogenicity) contain large amounts of immunogenic peptides (has high values of predicted T-cell immunogenicity scores).

### Descriptors calculation

Sequence based descriptors were primarily calculated using BioPython. Each Fv sequence pair was modelled using ABodyBuilder3 to obtain the structural information. The structure was then converted to PQR file using PDB2PQR (Dolinsky et al. 2004), with protonations and deprotonations predictions inferred by PROPKA3 (Olsson et al. 2011) at pH of 7.4. Using the PQR file, electrostatic potentials were calculated using the APBS (Jurrus et al. 2018) around the structure environment. Protein surface mesh was generated using Nanoshaper (Decherchi and Rocchia 2013) and for each vertex on the mesh, the potential was identified using the Multivalue program from APBS. The potentials were further assigned to the triangles of the mesh by calculating the mean potential value of the vertices constituting the face.

Similarly to PEP-patch (Hoerschinger et al. 2023) we calculate integrals of positively and negatively charged patches on the protein surface, by summing all triangles surfaces multiplied by their assigned potential.

### Data availability & access

We make the dataset available to non-profit organizations for non-commercial purposes. Other organizations & usage types, are kindly asked to contact us via https://naturalantibody.com/contact-us/

The dataset is available here: https://naturalantibody.com/therapeutic-antibody-database/

